# Glucose-mediated expansion of a gut commensal bacterium promotes *Plasmodium* infection through alkalizing mosquito midgut

**DOI:** 10.1101/2020.02.27.967315

**Authors:** Mengfei Wang, Yanpeng An, Shengzhang Dong, Yuebiao Feng, Li Gao, Penghua Wang, George Dimopoulus, Huiru Tang, Jingwen Wang

## Abstract

**SUMMARY:** Dietary sugar is the major energy source for mosquitoes, but its influence on mosquitoes’ capability to transmit malaria parasite remains unclear. Here we show that *Plasmodium berghei* infection changes global metabolism of *Anopheles stephensi* with the most significant impact on glucose metabolism. Supplementation of glucose or trehalose (the main hemolymph sugar) to mosquito increases *Plasmodium* infection by alkalizing the mosquito midgut. The glucose/trehalose diets promote rapid growth of a commensal bacterium, *Asaia bogorensis*, which remodels glucose metabolism and consequently increases midgut pH. The pH increase in turn promotes *Plasmodium* gametogenesis. We also demonstrate the sugar composition from different natural plants influences *A. bogorensis* growth and *Plasmodium* infection is associated with their capability to expand *A. bogorensis*. Altogether, our results demonstrate that dietary glucose is an important factor that determines mosquito’s competency to transmit *Plasmodium* and further highlight a key role for mosquito-microbiota metabolic interactions in regulating development of malaria parasite.

## INTRODUCTION

In addition to feed on animal blood, female mosquito also needs floral nectar and extra-floral plant fluids for energy provision. Sugar is the primary component of many floral nectars and a vital factor that determines mosquito fitness (Impoinvil et al., 2004; Magnarelli, 1978; Nayar and Sauerman Jr, 1971; Nayar and Sauerman, 1975). Sucrose and other sugars including glucose, mannose, galactose, fructose and gulose are commonly present in the nectar (Foster, 1995; Manda et al., 2007a; Manda et al., 2007b). The concentration of glucose, fructose, and gulose is correlated with mosquito’s survival and fecundity (Manda et al., 2007b). However, the influence of sugar metabolism on malarial parasite infection in mosquito remains poorly understood. As an obligate intracellular parasite, *Plasmodium* can sense nutrient levels of mammalian host and obtain them from host to fuel their growth and survival (Mancio-Silva et al., 2017). Glucose is the primary energy source for the extensive proliferation of malarial parasites at both the liver and blood stages (Kirk, 2001; Olszewski and Llinas, 2011). *Plasmodium* infection in hepatic cells leads to a significant increase in glucose uptake (Meireles et al., 2016). Erythrocytes infected with *Plasmodium* utilize glucose up to 100 times faster than uninfected cells (McKee et al., 1946; Meireles et al., 2016; Roth, 1987). Inhibition of glucose uptake of either parasite stage impairs its infection and attenuates virulence (Itani et al., 2014; Joët et al., 2003; Meireles et al., 2016; Saliba et al., 2004; Slavic et al., 2011). In *Anopheles* mosquito, indirect evidence shows that sugar also influences its susceptibility to transmit *Plasmodium*. Trehalose, composing of two molecules of glucose, is the major hemolymph (blood) sugar and is utilized after catabolized into glucose in insects (Becker et al., 1996). A reduction of trehalose level by knocking down trehalose transporter, AgTret1, impaired *Plasmodium falciparum* infection in mosquitoes (Liu et al., 2013). Insulin/insulin-like growth factor signaling (IIS) is also involved in regulating of *Plasmodium falciparum* infection in mosquitoes (Corby-Harris et al., 2010; Luckhart et al., 2013; Pietri et al., 2016). Thus, glucose metabolism in mosquito is expected to play a role in the development of *Plasmodium* parasites. However, the underlying mechanism remains unclear.

Mosquito is colonized by a vast and diverse consortium of microbes that are critical for nutrient assimilation. Eliminating gut microbiota by antibiotics delays food digestion (Gaio et al., 2011). *Serratia* produces hemolysins that might contribute to blood digestion in *Anopheles* mosquito (Chen et al., 2017b). *Acinetobacter* isolated from *Aedes albopictus* is able to digest blood containing α-keto-valeric acid and glycine, and nectar containing 4-hydroxy-benzoic acid and xylose (Minard et al., 2013). *Asaia bogorensis* isolated from *Anopheles* mosquitoes has a functional nitrogenase and might be responsible for the nitrogen metabolism in mosquitoes (Samaddar et al., 2011). In addition, gut microbiota also influence the outcome of pathogen infection through directly secreting anti-pathogen molecules and stimulating the basal immune responses (Bahia et al., 2014; Cirimotich et al., 2011a; Gonzalez-Ceron et al., 2003; Pumpuni et al., 1996). However, little is known about the sugar metabolic interactions between *Anopheles* and gut microbiota or the influence of such interactions on the capacity of mosquito to transmit pathogens.

In this study, we show that *Plasmodium* infection increases glucose consumption in *An. stephensi* by metabolomics analyses. Oral administration of glucose and trehalose promotes *Plasmodium* infection, respectively. Such an effect relies on the presence of gut microbiota. Glucose supplementation enriches a gut commensal bacterium, *A. bogorensis*, which changes glucose metabolism and increases pH of mosquito midgut. The alkalization of *An. stephensi* midgut promotes *P. berghei* sexual development. We further explore the potential correlation between the abundance of *A. bogorensis* and the capacity of *An. stephensi* to transmit *Plasmodium* by providing mosquito with sugar meals simulating the sugar composition of plants in malaria endemic area. Collectively, our results demonstrate that *A. bogorensis* participates the glucose metabolism in *An. stephensi* and such interplay determines the outcome of parasite infections.

## RESULTS

### *P. berghei* infection reduces glucose and trehalose levels in *An. stephensi*

To determine whether *Plasmodium* infection influences mosquito metabolism, we analyzed the metabolomics changes of whole *An. stephensi* 1 day post *P. berghei* infection (1 dpi) using nuclear magnetic resonance spectroscopy (NMR) analysis. This is the stage when ookinetes invade midgut epithelium and elicit a global change of gene expression (Domingos et al., 2017; Dong et al., 2006). *P. berghei* infection induced profound metabolic changes in *An. stephensi*. Totally 21 metabolites were differentially regulated (Figure 1 and Table S1). Among them, the metabolites associated with glucose metabolism including trehalose, glucose, succinate, citrate were significantly reduced 1 dpi, while pyruvate and acetate were significantly increased (Figure 1B). These data suggest that *Plasmodium* infection disrupts the homeostasis of sugar metabolism in *An. stephensi*.

**Figure 1.**
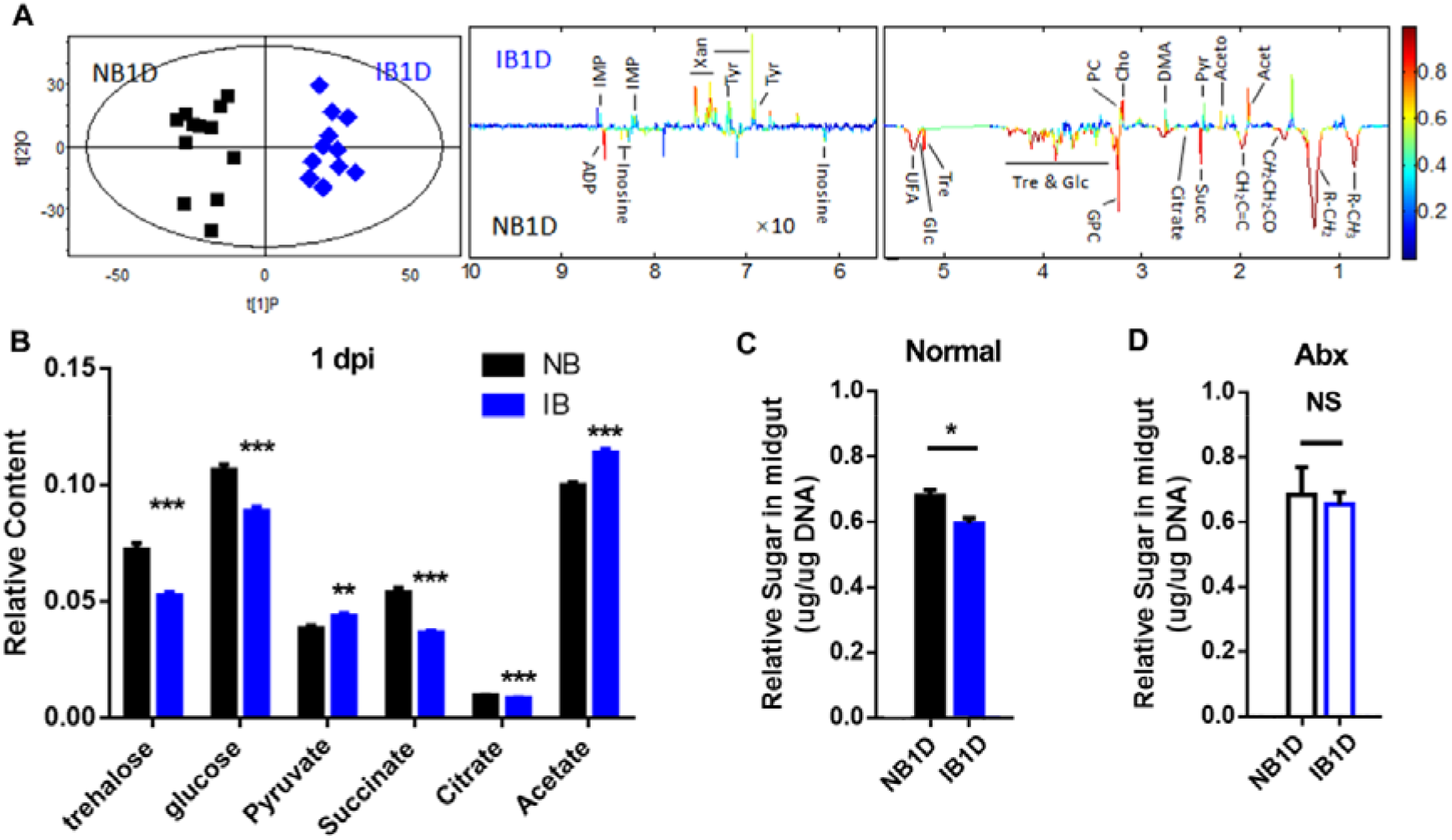
*P. berghei* infection changes glucose metabolism in *An. stephensi*. (A)OPLS-DA scores plots (left) and corresponding loadings plots (right) showed the metabolites significantly changed in the mosquito extract fed on normal blood (NB) and infectious blood (IB) 1 day post (1 dpi) infection. R^2^X=0.430, Q^2^=0.857. Metabolite keys are the same as in Table S1. (B) Relative content of the key sugar metabolites. All of the integral regions were normalized to dried weight of mosquitoes for each spectrum, respectively. (C and D)The relative concentration of total sugar level (glucose and trehalose) in the midgut of mosquito treated without (C) or with antibiotics (Abx) (D) 1 dpi. Glucose and trehalose concentrations were normalized to genomic DNA extracted from midgut. Significance in A was determined by CVANOVA approach with p < 0.05 as the significant level. Significance in B-D was determined by Student’s *t* test. Error bars indicate standard error of the mean (*n* = 5). NS, no significance, *, p <0.05, **, p<0.01, and ***, p<0.001.

We next specifically examined the total glucose and trehalose level in the midgut because midgut is the tissue where *Plasmodium* first invades and undergoes gametogenesis, fertilization and ookinete formation (Angrisano et al., 2012). *P. berghei* infection significantly decreased total sugar level in the midgut (Figure 1C). Because mosquito midgut is in constant association with commensal bacteria, to determine whether gut microbiota is also involved in sugar reduction during *Plasmodium* infection, we analyzed the total glucose and trehalose level in midguts of mosquitoes treated with a cocktail of antibiotics. Interestingly, when the majority of gut microbiota was removed, *P. berghei* infection failed to decrease the total glucose and trehalose level in midgut (Figure 1D). These results indicate that gut microbiota contributes to the *Plasmodium*-mediated alteration of glucose metabolism in mosquitoes.

### Increasing uptake of glucose/trehalose promotes *Plasmodium* infection in *An. stephensi*

To examine the influence of glucose and trehalose on *Plasmodium* infection, we first orally supplemented 2% sucrose containing glucose (G), trehalose (T), or 10% sucrose (HS) to newly emerged mosquitoes for 5 days, respectively, followed by infecting mosquitoes with *P. berghei* and *P. falciparum* (Figure 2A). The number of *P. berghei* oocysts increased significantly with addition of 0.1 M glucose and trehalose, respectively, compared to sucrose- only controls (S and HS) (Figure 2B&S1A). Trehalose (0.1 M) similarly increased oocyst intensity of *P. falciparum* compared to sucrose controls (S) (Figure 2C). However, simply increasing sucrose concentration (10% sucrose, HS) failed to promote infection of either *P. berghei* or *P. falciparum* (Figure 2B&C). Oral administration of glucose (G) diet to mosquitoes post infectious blood meal had no influence on parasite infection (Figure S1B), suggesting that certain alteration resulted from glucose/trehalose supplementation before *Plasmodium* infection contributes to the elevated susceptibility.

**Figure 2.**
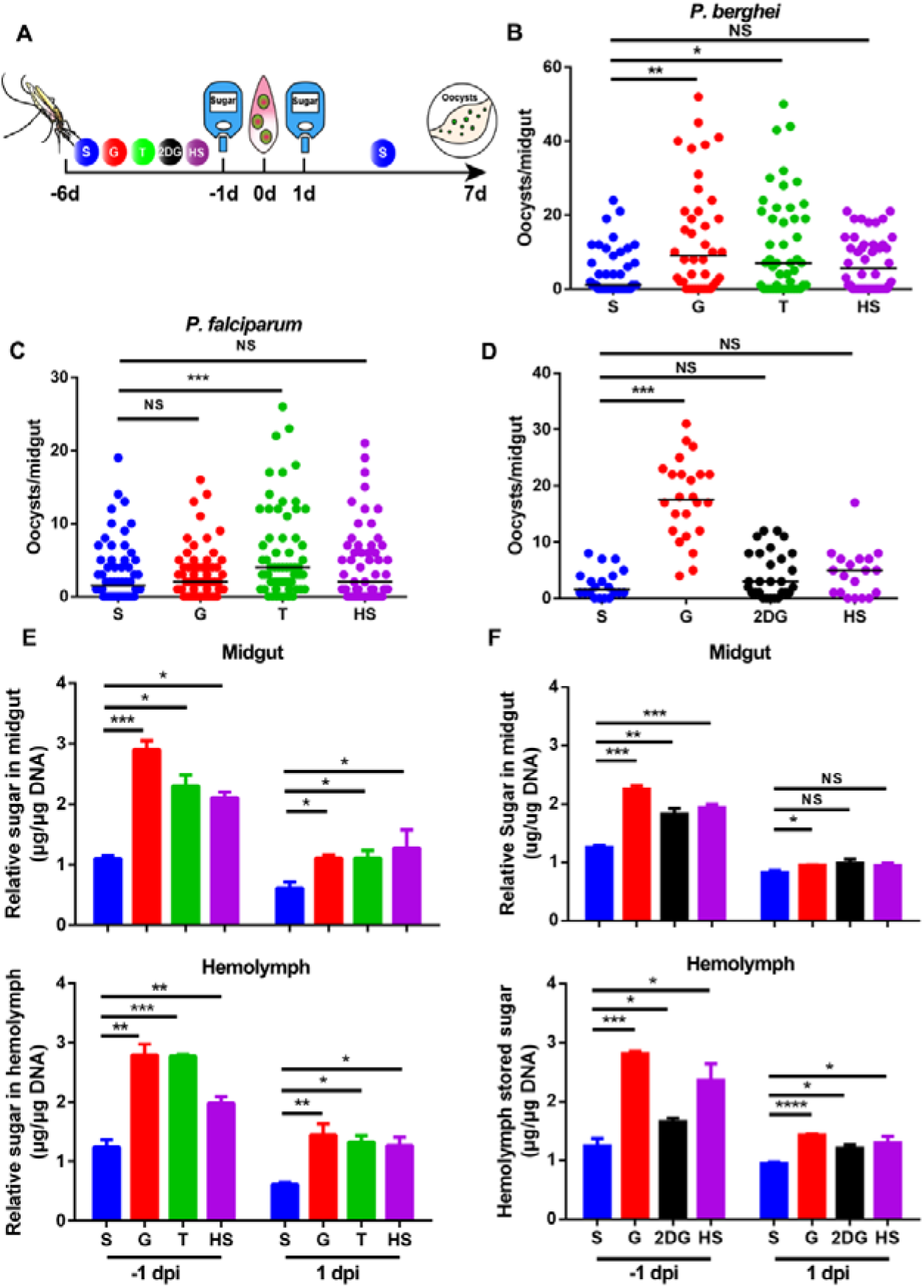
Glucose and trehalose supplementation promotes *Plasmodium* infection in *An. stephensi*. (A) Work flow of sugar treatment on *Anopheles* mosquito. (B and C) *P. berghei* and *P. falciparum* oocyst intensities in mosquito fed with S, G, T and HS diets, respectively. (D) *P. berghei* oocyst intensity in mosquitoes fed with S, G, 2-DG and HS diets. (E and F) The relative concentration of total sugar (glucose and trehalose) in the midgut (top panel) and hemolymph (bottom panel) of mosquitoes fed on the sugar diets in correspondence to (B) and (D), respectively, 1 day prior to (−1 dpi) and 1 day (1 dpi) post infection. Glucose and trehalose concentration was normalized to the amount of genomic DNA extracted from midgut and hemolymph, respectively. Each dot represents an individual mosquito and horizontal lines represent the medians. Results shown in B-D were pooled from at least 2 independent experiments. Significance was determined by Mann-Whitney test. Significance in E and F was determined by Student’s *t* test. Error bars indicate standard error of the mean (*n* = 5). NS, no significance, *, p <0.05, **, p<0.01, ***, p<0.001 and ****, p<0.0001. S, 2% sucrose; G, 2% sucrose+0.1 M glucose; T, 2% sucose+0.1 M trehalose; 2-DG, 2% sucrose+0.1 M glucose+5 mM 2-DG; HS, 10% sucrose.

*P. berghei* infection decreased glucose, while increased pyruvate, suggesting that parasite invasion might increase glycolysis activity. To examine the increasing susceptibility of glucose-supplemented mosquitoes to parasite infection is due to the elevated glucose or the enhanced glycolysis, we fed mosquitoes with two glycolysis intermediates, glucose-6-phosphate, frucotose-1, 6-bisphosphate and one product, pyruvate. None of these metabolites influenced parasite infection in *An. stephensi* (Figure S1C), suggesting that the increased *Plasmodium* infection was due to the supplementation of glucose. We next introduced glycolysis inhibitor 2-deoxy-D-glucose (2-DG) in combination with glucose to mosquitoes before *Plasmodium* infection. Supplementation of 2-DG completely restored the susceptibility to *P. berghei* infection to control level (Figure 2D). These results suggest that increasing glucose and trehalose uptake in *An. stephensi* before parasite infection promotes *Plasmodium* parasites infection.

*Plasmodium* rely on host glucose for energy needs (Olszewski and Llinas, 2011), we next examined whether glucose-mediated enhancement in *P. berghei* infection could be due to the increasing availability of sugar in *An. stephensi*. The total trehalose+glucose level was examined in midguts and hemolymph 1 day prior to (−1 dpi) and 1 day post (1 dpi) infection, respectively (Figure 2E). Increasing sugar uptake including feeding mosquito with G, T, and HS diet all significantly increased sugar level in midguts and hemolymph −1 dpi and 1dpi (Figure 2E). Treatment with 2-DG also accumulated total sugar level significantly comparing to controls (S) in both midguts and hemolymph before *Plasmodium* infection (Figure 2F). However, the accumulation of sugar level in HS- and 2-DG-fed mosquitoes failed to increase oocyst load. Altogether, these data show that glucose and trehalose-mediated increase in susceptibility of *An. stephensi* to *Plasmodium* infection is not due to the elevated availability of glucose and trehalose in mosquitoes.

### *Asaia bogorensis* expanded by glucose/trehalose supplementation facilitates *P. berghei* infection in *An. stephensi*

Gut microbiota is an important factor that determines vector competence (Ricci et al., 2012). In combination with our finding that *P. berghei*-induced reduction of total sugar relies on the presence of gut microbiota, we first examined whether gut microbiota was involved in glucose-mediated increasing susceptibility to parasite infection. Mosquitoes were fed with S and G containing a cocktail of antibiotics to eliminate gut microbiota followed by feeding on *P. berghei*-infected mice. The oocyst number was counted 7 days post infection (Figure 3A). Glucose supplementation significantly increased the number of *P. berghei* oocysts in normal mosquitoes, but failed to do so in antibiotics-treated ones (Figure 3A). *An. stephensi* deprived of microbiota had significantly higher sugar levels than that in normal ones, further confirming that gut microbiota is responsible for the sugar consumption (Figure S2A).

**Figure 3.**
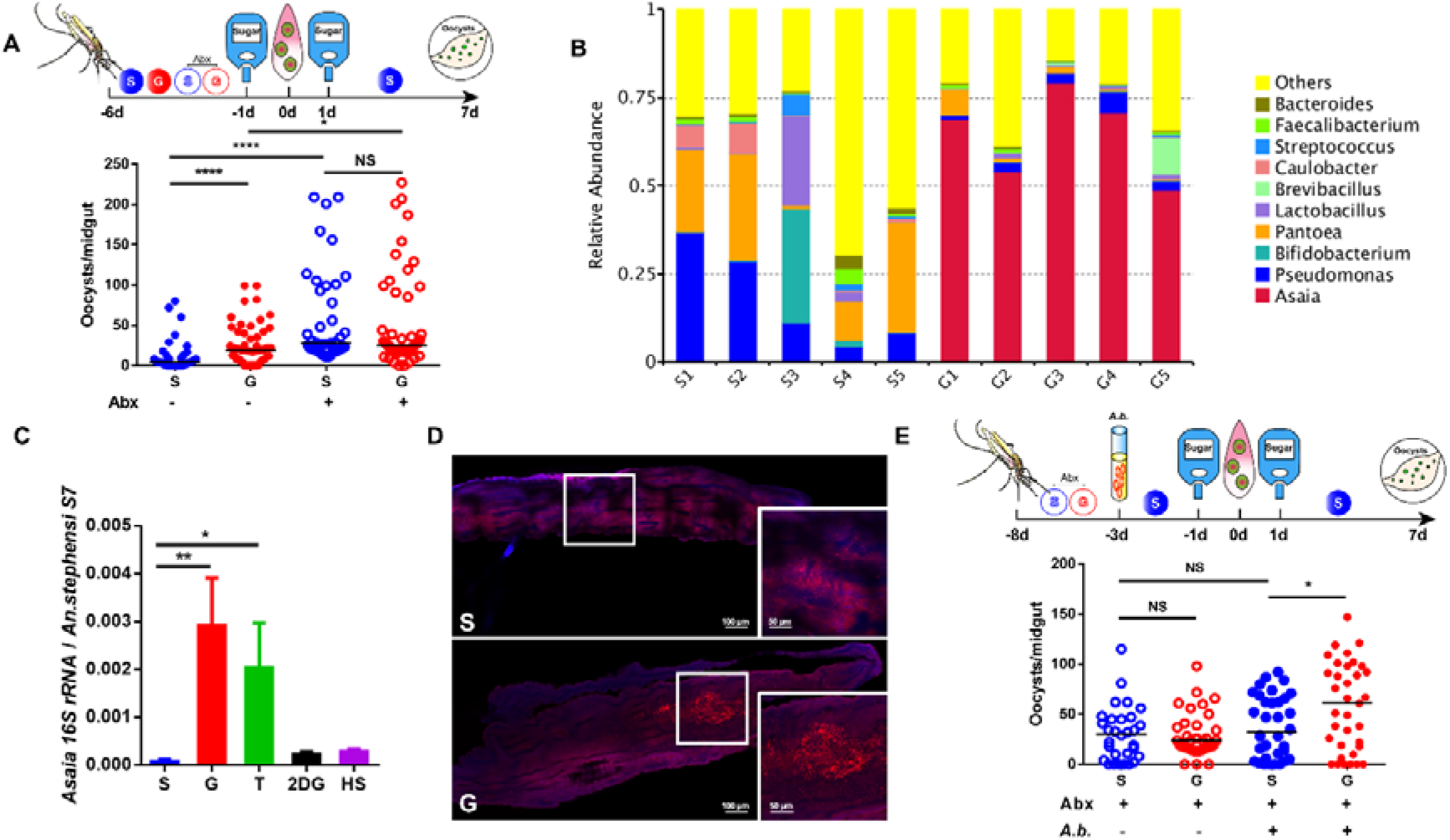
*Asaia bogorensis* promotes *P. berghei* infection in *An. stephensi*. (A) Influence of antibiotics treatment on *P. berghei* infection. (B) Relative abundance of major bacteria genera in *An. stephensi* midguts fed on S and G by 16S rRNA pyrosequencing. Each column represents 3 pooled midguts. (C) The abundance of *A. bogorensis* in the midgut of *An. stephensi* determined by qPCR. (D) Localization of *A. bogorensis* (red) in the midgut of *An. stephensi* on S and G, using *A. bogorensis* specific probes. Nuclei were stained with DAPI (blue). The enlarged view in the box is shown on the right of the figure. Images are representative of at least 5 individual mosquito midguts. (E) Influence of *A.bogorensis* re-colonization on *P. berghei* infection outcome. Each dot represents an individual mosquito and horizontal lines represent the medians. Results shown in A and E were pooled from 2 independent experiments. Significance was determined by Mann-Whitney test. Significance in C was determined by Student’s *t* test. Error bars indicate standard error of the mean (*n* = 12). NS, no significance, *, p <0.05, **, p<0.01, and ****, p<0.0001. S, 2% sucrose; G, 2% sucrose + 0.1 M glucose; Abx, antibiotics treatment; *A.b.*, *A. bogorensis*.

To search for the specific bacteria responsible for sugar catabolism, we analyzed the community structure of gut microbiota of mosquitoes feeding on S and G by 16S rRNA pyrosequencing, respectively (Figure 3B and Figure S2B&C). A total 729,898 reads with a median length of 415.9 bp were obtained, identifying 1,879 bacterial operational taxonomic units (OTUs). Principal component analysis showed that gut microbiota of mosquitoes fed on G diet clustered separately from those on S controls (Figure S2B). Glucose supplementation significantly increased the relative abundance of *Asaia sp.* in *An. stephensi* midgut compared to S group (Figure 3B and Figure S2C) and *Asaia bogorensis* was the bacterial species enriched by glucose diet in *An. stephensi*. The result of real-time quantitative PCR confirmed that supplementation of glucose and trehalose significantly increased the number of *A. bogorensis*, respectively (Figure 3C). Increased fluorescent signals were also observed in midguts of *An. stephesi* on G diet compared to control by fluorescent in situ hybridization (FISH) analysis (Figure 3D). These findings suggest that addition of glucose and trehalose promotes proliferation of *As. bogorensis*. This bacterium might participate in the glucose- and trehalose-mediated enhancement in *P. berghei* infection.

We next colonized antibiotics-treated mosquitoes with *A. bogorensis* and analyzed susceptibility of these mosquitoes to *P. berghei* infection. Two days post-inoculation, *A. bogorensis* colonized the midgut successfully as the abundance of this bacterium was significantly higher than that in normal mosquitoes (Figure S2D). Re-colonization of *A. bogorensis* alone was not able to change infection rate in comparison to antibiotics-treated ones (Figure 3E). However, addition of glucose significantly increased median oocyst number from 32 to 61 in *A. bogorensis* mono-associated mosquitoes (Figure 3E). Moreover, colonization of *A. bogorensis* significantly reduced sugar level in the mosquito midgut (Figure S2E). Altogether, these data indicate that glucose and trehalose administration to mosquitoes promotes *A. bogorensis* proliferation in the midgut. The expansion of *A. bogorensis* renders mosquito more permissive to parasite infection.

### Glucose/trehalose promotes *P. berghei* infection by disturbing midgut acid-base balance

To determine how glucose promotes *P. berghei* infection, we first performed an RNA-sequencing (RNAseq) analysis on *An. stephensi* midguts fed on S and G −1 dpi and 1 dpi, respectively. A total 460 and 97 genes were differentially expressed −1 dpi and 1 dpi, respectively (Table S2). Supplementation of glucose significantly up-regulated the expression of genes associated with glucose and other sugar metabolism, and sugar transportation −1 dpi. However, most of these genes were down-regulated 1 dpi (Figure 4A). Notably, glucose and trehalose supplementation significantly repressed the activity of Imd pathway (Figure 4A and Figure S3A). The expression level of *caudal*, the negative regulator of IMD pathway was significantly up-regulated, while peptidoglycan recognition proteins, *pgrp-lc*, antimicrobial peptides, *defensin*, and *cecropin* were significantly down-regulated both prior to and post infectious blood meal (Cirimotich et al., 2010; Clayton et al., 2013; Clayton et al., 2014; Song et al., 2018)(Figure 4A). It is highly possible that increased susceptibility to *P. berghei* infection results from the down-regulation of Imd pathway. We next stimulated the expression of Rel2-mediated immune genes by knocking down *caudal*. However, supplementing glucose to these mosquitoes similarly increased their oocysts number compared to those on S diet (Figure S3B). Neither knocking down of *caudal* affected the abundance of *A. bogorensis* (Figure S3C). Thus, Imd pathway is not responsible for glucose-mediated enhancement in *P. berghei* infection (Garver et al., 2009).

**Figure 4.**
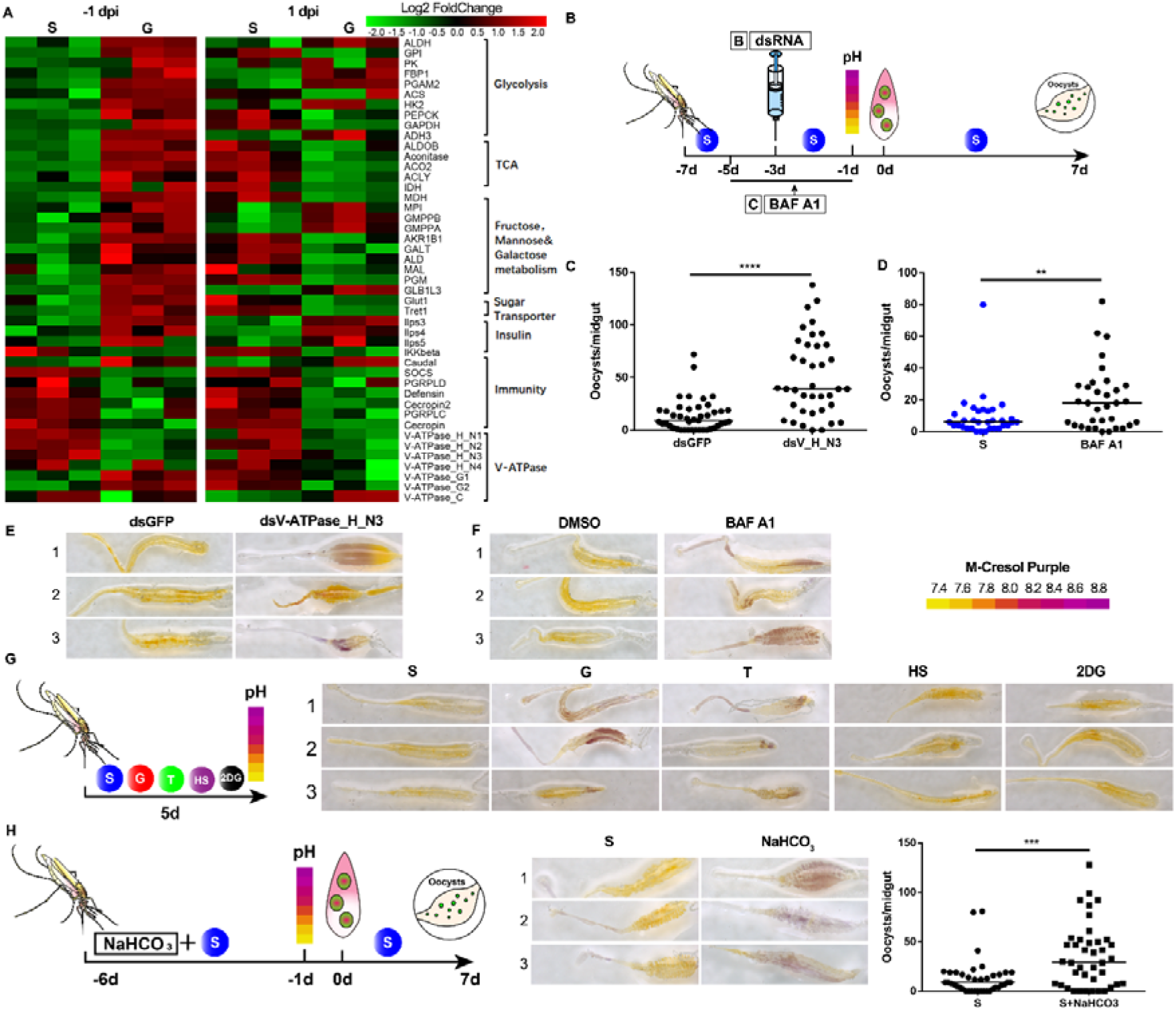
Midgut pH determines the susceptibility of *An. stephensi* to *P. berghei* infection. (A) Hierarchical clustering analysis of differentially expressed genes in mosquito midguts fed with S and G, respectively, −1 dpi and 1dpi. Upregulated genes are shown in red; downregulated genes are shown in green. (B) Work flow of dsRNA injection (C) and BAF A1 treatment (D) in *An. stephensi*. (C and D) *P. berghei* oocyst number in dsRNA (C) and BAF A1 (D) treatments. (E and F) The pH staining of mosquito midguts treated with dsRNA (E) and BAF A1 (F) by pH indicator m-cresol purple. The pH indicator is shown on the right. Images are the three representatives of at least 5 individual mosquito midguts. (G) The pH staining of mosquito midguts fed on S, G, T, HS and 2-DG diets by m-cresol purple, respectively. Work flow of pH assay of mosquitoes on S, G, T, HS and 2-DG diets (left). The pH staining of midguts on different diets (right panel). Images are the three representatives of at least 5 individual mosquito midguts. (H) The effect of NaCHO_3_ supplementation on *P. berghei* infection. Work flow of NaCHO_3_ treatment and pH assay (right panel), pH staining of NaHCO_3_ feeding midguts (middle panel), *Plasmodium* oocyst numbers after NaHCO_3_ treatment (right panel). Images are the three representatives of at least 5 individual mosquito midguts. Each dot represents an individual mosquito and horizontal lines represent the medians. Data shown in C, D and H were pooled from 2 independent experiments. Significance was determined by Mann-Whitney test. NS, no significance, *, p <0.05, **, p<0.01, ***, p<0.001 and ****, p<0.0001. S, 2% sucrose; G, 2% sucrose+0.1 M glucose; T, 2% sucose+0.1 M trehalose; 2-DG, 2% sucrose+0.1 M glucose+5 mM 2-DG; HS, 10% sucrose, NaCHO_3_, 2% sucrose+ 0.1 M NaCHO_3_.

Moreover, we observed a group of genes encoding Vacuolar ATPase (V-ATPase), including vacuolar-type H(+)-ATPase*(V-ATPase)*_*H*_*N1*, _*H*_*N2*, _*H*_*N3*, _*H*_*N4*, _*G1*, _*G2* and _*C*, were differentially regulated when *An. stephensi* was fed with G −1 dpi and 1 dpi, respectively (Figure 4A). V-ATPase hydrolyzes ATP and transports proton across membranes. It is critical for controlling the intracellular and extracellular pH of cells (Boudko et al., 2001a; Hayek et al., 2019; Linser et al., 2009). To examine whether V-ATPase is responsible for the glucose-induced increase of vector competence, we impaired V-ATPase function by specifically knocking down a *V-ATPase* gene encoding cytoplasmic V1 domain subunit H (*V-ATPase_H_N3*) or supplementation of a V-ATPase inhibitor, Bafilomycin A1, in normal S feeding mosquitoes (Figure 4B). Silencing this gene significantly increased oocyst number compared to dsGFP controls in mosquitoes feeding on S (Figure 4C and S3D). Again, orally administration of Bafilomycin A1 to *An. stephensi* dramatically increased oocyst number of *P. berghei* compared to controls (Figure 4D) (Boudko et al., 2001b; Yoshimori et al., 1991). These results indicate that glucose-induced enhancement in *P. berghei* infection may be a result of dysfunctional V-ATPase.

As V-ATPase is involved in maintaining acid base balance in mosquito midguts (Jagadeshwaran et al., 2010; Patrick et al., 2006; Zhuang et al., 1999), we next examined whether disruption of V-ATPase expression could influence midgut pH. As expected, inhibition of V-ATPase by dsV-ATPase treatment and Bafilomycin A1 supplementation both increased midgut pH compared to controls by *m*-Cresol purple pH staining assay (Figure 4E&F). Thus, these results indicate that inhibiting V-ATPase activity led to an increase in midgut pH.

As glucose regulated V-ATPase expression and V-ATPase determined midgut pH, we next examined whether glucose supplementation could influence midgut pH. Mosquitoes were fed with S, G, T, HS and 2-DG for 5 days and midguts were dissected for pH analysis (Figure 4G). Addition of glucose and trehaose increased midgut pH remarkably compared to controls, however, increasing sucrose concentrations didn’t change midgut pH (Figure 4G). Again, inhibition of glycolysis by 2-DG restored glucose-induced pH back to the control level (Figure 4G). Thus, our results show that administration of glucose and trehalose dysregulates expression of V-ATPase that is responsible for maintaining homeostasis of midgut pH.

To examine whether elevated pH enhances *Plasmodium* infection, we next feeding *An. stephensi* with 2% S containing different concentrations of NaHCO_3_ (Figure S3E). Addition of 0.1 M NaHCO_3_ dramatically increased midgut pH and significantly increased oocyst numbers compared to that in controls (Figure 4H and S3F). In these mosquitoes, we also observed a significant reduction in Imd signaling activity both before and after *P. berghei* infection, suggesting that the downregulation of immune activity might be a result of pH increase (Figure S3G). Taken together, these findings suggest that glucose/trehalose administration in mosquitoes disturbs pH homeostasis in the midgut, which facilitates *Plasmodium* infection.

### *A. bogorensis* is responsible for glucose/trehalose-mediated alkalization of mosquito midgut

In light of our finding that glucose and trehalose supplementation enriches gut commensal *A. bogorensis* and increases midgut pH, we were interested to know whether *A. bogorensis* could be responsible for controlling midgut acid-base balance. The midgut pH of antibiotics-treated mosquitoes mono-colonized by *A. bogorensis* fed on S and G was analyzed by *m*-Cresol purple dye, respectively. Antibiotics-treated mosquitoes on the same diet were used as controls (Figure 5A). Neither glucose diet nor re-colonization of *A. bogorensis* alone alkalized mosquito midgut (Figure. 5A). However, re-colonization of *A. bogorensis* in antibiotics treated mosquitoes fed with G increased midgut pH dramatically compared to the ones with the bacterium alone (Figure 5A). In combination with our previous finding that glucose selectively expands *A. bogorensis*, these results indicate that glucose catabolic activity of *A. bogorensis* might contribute to the pH changes in mosquito midgut.

**Figure 5.**
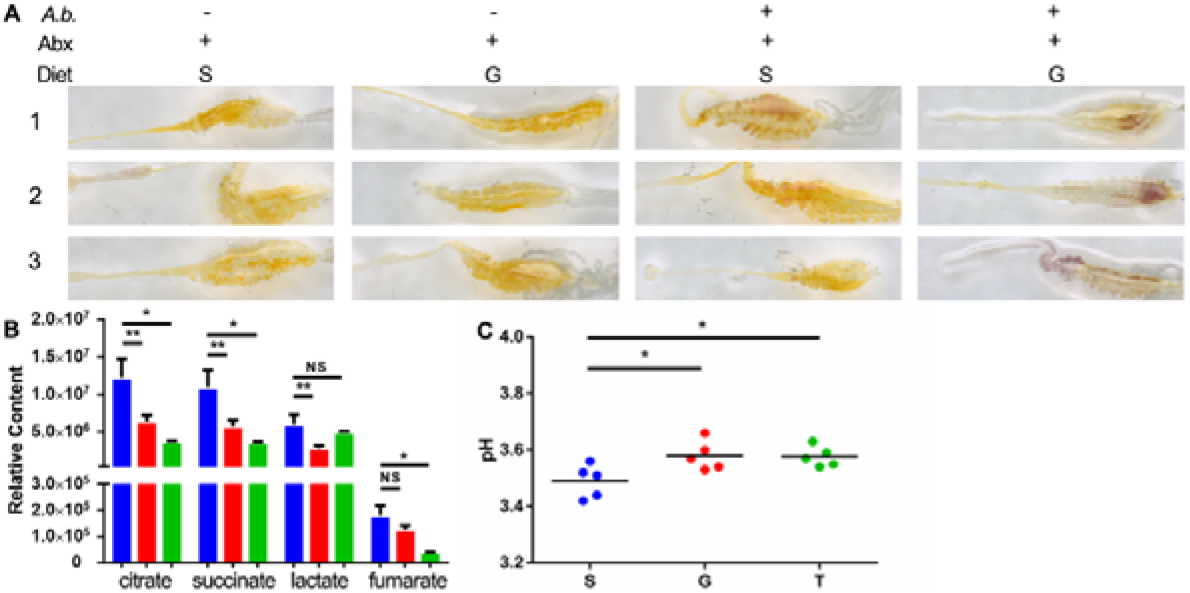
Glucose metabolic activity of *A. bogorensis* determines the midgut pH. (A) Influence of *A.bogorensis* re-colonization on the pH of mosquito midgut. Images are the three representatives of at least 5 individual mosquito midguts. (B) Relative content of the key sugar metabolites in the three conditioned mediums. (C) The pH value of S (blue), G (red) and T (green) containing medium 3 days post *A. bogorensis* growth. Significance in B and C was determined by Student’s *t* test. Error bars indicate standard error of the mean (*n* = 5). NS, not significant, *, p <0.05, and **, p<0.01. S, sucrose; G, glucose; T, trehalose.

As we have technical difficulties in analyzing metabolites of *A. bogorensis* in the mosquito migut, we next compared the difference of sugar metabolites in conditioned media from *A. bogorensis* grown in the presence of 0.1 M sucrose, glucose and trehalose for 1 week, respectively, by NMR analysis. Use of glucose as a carbon source significantly altered the level of 13 metabolites with 3 glucose metabolites, citrate, succinate, and lactate significantly reduced (Figure 5B and Table S3). The trehalose-containing medium also decreased the levels of citrate, succinate, and fumarate significantly (Figure 5B). The reduction of multiple acids might lead to the increased pH of the medium. We next examined pH of the conditioned medium and found the pH was higher in glucose and trehalose-containing medium than that in sucrose-containing medium, respectively (Figure 5C). These data suggest that the proliferation of *A. bogorensis* changes glucose metabolism, which in turn increases midgut pH.

### Alkalization of mosquito midgut promotes *P. berghei* gametogenesis

An increase of pH from mammalian host (pH=7.2) to mosquito midgut (pH=8) is a key trigger for *Plasmodium* male gametogenesis (Bennink et al., 2016; Billker et al., 1997; Carter and Nijhout, 1977; Kawamoto et al., 1991). Based on our finding that expansion of *A. bogorensis* increased midgut pH, we hypothesized that such pH elevation could benefit *P. berghei* sexual development in the midgut. Gamete is developed around 15 min after *Plasmodium* is ingested into mosquito midgut (Bennink et al., 2016). We first quantified the expression of 6 genes associated with *Plasmodium* male gametogenesis in the midgut of *An. stephensi* fed on the four sugar diets, S, G, 2-DG and HS 10 min post infection (Figure 6A). They are calcium-dependent protein kinase 1 (*cdpk1*), protein phosphatase 1 (*ppm1*), gamete egress and sporozoite traversal protein (*gest*), male development gene-1 (*mdv-1*), basal body protein SAS-6 (*sas-6*), the sexual stage-specific actin isoform actin 2 (*actin 2*)(Billker et al., 2004; Deligianni et al., 2011; Guttery et al., 2014; Lanfrancotti et al., 2007; Marques et al., 2015; Olivieri et al., 2015; Ponzi et al., 2009; Sebastian et al., 2012; Sturm et al., 2015; Talman et al., 2011; Theriot et al., 2014; Yeoh et al., 2017; Yuda et al., 2015). As expected, 5 of 6 genes including *cdpk1*, *ppm1*, *sas-6*, *gest*, and *mdv-1* were significantly upregulated in glucose-fed midguts. While feeding on either 2-DG or HS didn’t influence the expression level of most of these genes (Figure 6A). We next analyzed the expression level of the same 6 genes in midguts of *A. bogorensis* re-colonized *An. stephensi* fed on S and G 10 min post infection (Figure 6B). Colonization of *A. bogorensis* in glucose-fed mosquitoes significantly elevated expression of 4 of 6 genes associated with microgamete development (Figure 6B). Similarly, addition of NaCHO_3_ induced the expression of most of these genes (Figure S4). To further confirm that increase of pH promotes male gametogenesis, we visualized the exflagellation by staining the blood bolus from mosquito midguts using Giemsa staining. There was a significant higher number of exflagellations and exflagellation rate in G and T fed-mosquitoes, respectively, than that in S fed-ones (Figure 6C&D). These results suggest that elevated midgut pH upregulated expression of gametogenesis-related genes, thereby promoting male gamete formation in the midgut.

**Figure 6.**
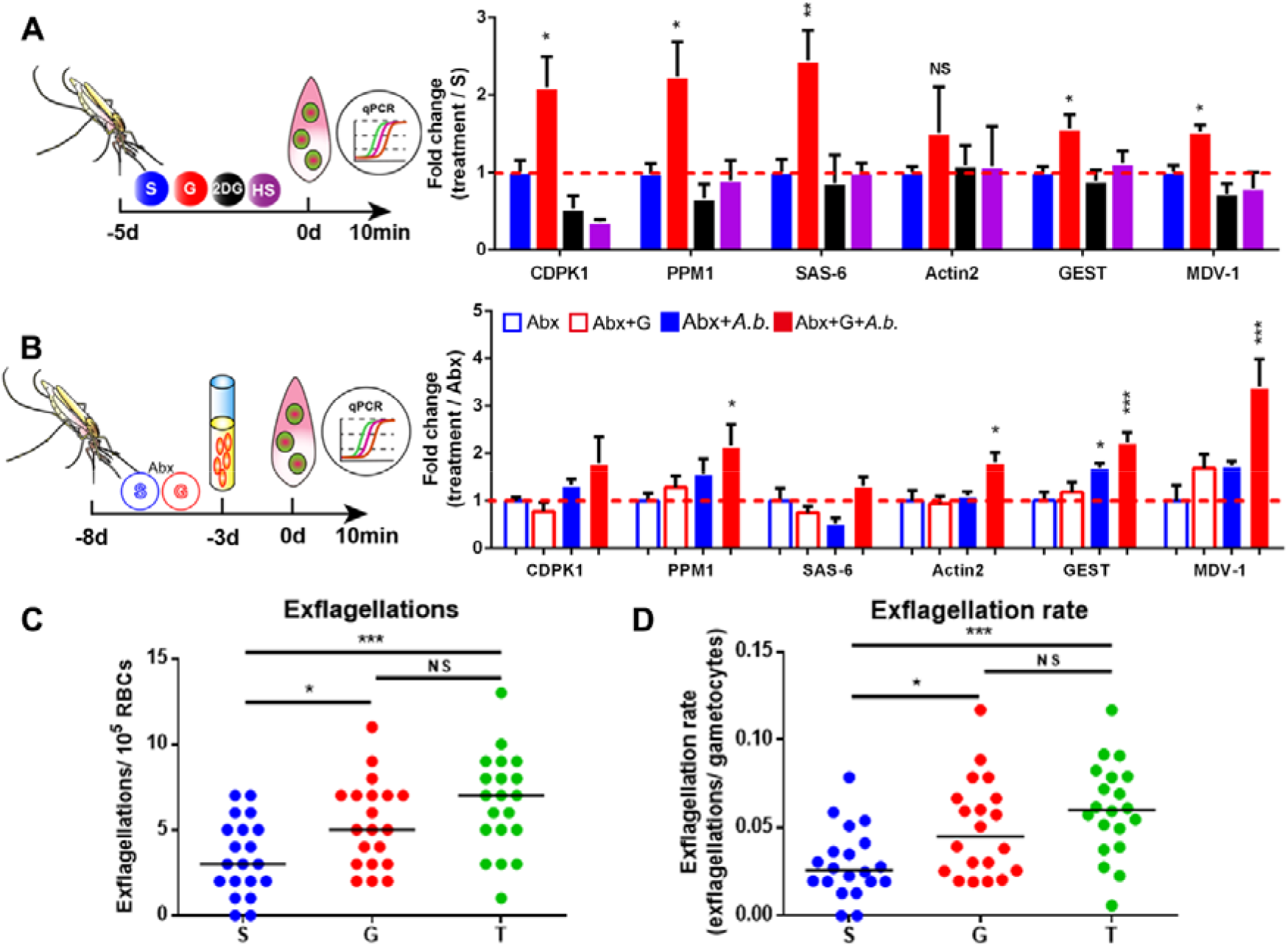
The increase of pH induces male gametogenesis. (A) Quantification of the expression level of genes associated with male gametogenesis in mosquitoes fed with S, G, 2-DG and HS, respectively. (B) Quantification of the expression level of genes associated with male gametogenesis in *A. bogorensis* re-colonized mosquitoes fed with S and G, respectively. (C and D) Exflagellations and exflagellation rate in the mosquitoes fed on S, G and T, respectively. Exflagellations were monitored 10 min post infection. The median number of exflagellations per 10^5^ erythrocytes is shown (C) and exflagellation rate was shown as the percent (exflagellations/gametocytes) per 10^5^ erythrocytes (D). Significance in A and B was determined by Student’s *t* test. Error bars indicate standard error of the mean (*n* = 5). Each dot represents one mosquito and horizontal lines represent the medians in C and D. Data shown in C and D was pooled from 2 independent experiments. Significance was determined by Mann-Whitney test. NS, no significance, *, p <0.05, **, p<0.01, and ***, p<0.001. S, 2% sucrose; G, 2% sucrose+0.1 M glucose; T, 2% sucrose+0.1 M trehalose; 2-DG, 2% sucrose+0.1 M glucose+5 mM 2-DG; HS, 10% sucrose.

### Sugar composition of natural plants influences the abundance of *A. bogorensis* and vector competence

The diversity of natural plants affects the capacity of *Anopheles* mosquito to transmit *Plasmodium* parasites (Ebrahimi et al., 2018; Gu et al., 2011; Hien and Dabire, 2016). We hypothesized that the ability of different sugar composition to promote the growth of *A. bogorensis* might vary and such variation might influence the vector competence of *Anopheles* mosquitoes. We compared the abundance of *A. bogorensis* in mosquitoes feeding on sugar meals simulating the nectar of 5 plants in western Kenya before blood meal. They are *Tecoma stans*, *Senna didymobotrya*, *Ricinus communis*, *Parthenium hysterophorus* and *Lantana camara* (Manda et al., 2007a)(Figure 7A). Flowers of *T. stans* and *S. didymobotrya*, and leaves and stems of *R. communis* and *P. hysterophorus* are the favorite resources of *An. gambiae* in western Kenya, while plant sap of *L. camara* leaves and stems is the least preferred (Manda et al., 2007a; Manda et al., 2007b). Sugar solution based on the composition and water content of five plant species were prepared (Abebe, 2015; Abere and Enoghama, 2015; Fatimah and Ahmad, 2009; Manda et al., 2007a; Sowmya et al., 2019). The *T. stans*, *S. didymobotrya* meals significantly increased the abundance of *A. bogorensis* as glucose did (Figure 7B). While, the sugar meals simulating *P. hysterophorus* and *R. communis* significantly decreased this bacterium level comparing to S group (Figure 7B). We next examined the susceptibility of these mosquito to *P. berghei* infection. As expected, mosquitoes in which *A. bogorensis* was expanded had significantly higher oocyst number compared to S controls (Figure 7C). However, the reduction of *A. bogorensis* in *P*. *hysterophorus* and *R. communis-*fed mosquitoes didn’t further inhibit parasite infection (Figure 7C). Although *T. stans*, *S. didymobotrya*, *R. communis* and *P*. *hysterophorus* are the four favorite plant species of *Anohpeles mosquitoes*, their ability to expand *A. bogorensis* varies. Those promote proliferation of *A. bogorensis* also facilitate *Plasmodium* infection in mosquitoes.

**Figure 7.**
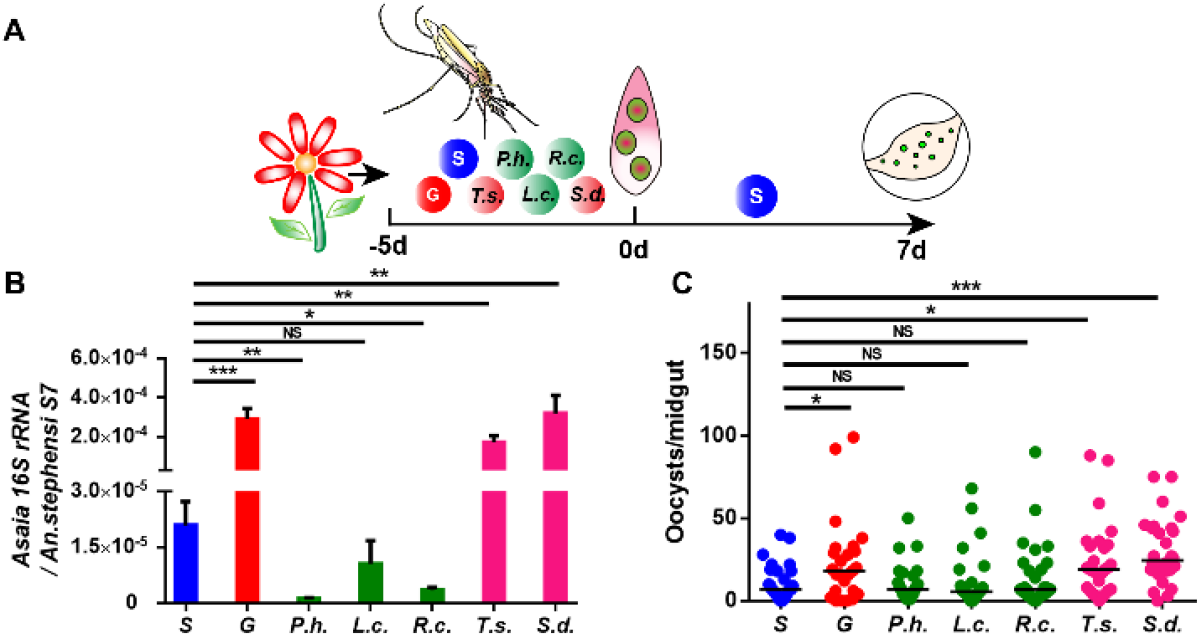
Sugar composition affects the abundance of *A. bogorensis* and *P. berghei* infection outcome. (A) Work flow of sugar treatment in *An. stephensi*. (B) Quantification of levels of *A. bogorensis* in mosquitoes fed with different sugar diets. (C) Oocyst number in mosquitoes fed on different sugar diets. Significance in B was determined by Student’s *t* test. Error bars indicate standard error of the mean (*n* = 12). Each dot represents an individual mosquito and horizontal lines represent the medians in C. Data shown in C was pooled from 2 independent experiments. Significance was determined by Mann-Whitney test. NS= no significance, *, p <0.05, **, p<0.01, and ***, p<0.001. S, 2% sucrose; G, 2% sucrose+0.1 M glucose; *P.h*., *P*. *hysterophorus*, *R.c.*, *R. communis*, *L.c.*, *L. camara*, *T.s.*, *T. stans*, *S.d.*, *S. didymobotrya*.

## DISCUSSION

Sugar is a major energy source for mosquitoes, but its influence on the capability of mosquito to transmit pathogens remains unclear. *Plasmodium* parasites have lost multiple genes involved in nutrient synthesis and have to compete with their hosts for essential nutrients (Gardner et al., 2002). Both liver and blood stage malaria parasites scavenge host glucose for their energetic needs (Meireles et al., 2016; Olszewski and Llinas, 2011; Stanway et al., 2019; Wang et al., 2018). However, little is known about the crosstalk between glucose metabolism of *Anopheles* and that of *Plasmodium*. In this study we demonstrate that *Plasmodium* infection disrupts the homeostasis of glucose metabolism in *An. stephensi*. Such disturbance relies on the presence of commensal bacterium *A. bogorensis*. The expansion of this bacterium changes glucose metabolism and increases pH in the midgut, thereby promoting *Plasmodium* gametogenesis and infection.

Different from the rapid proliferation in mammalian stage, *Plasmodium* undergoes a drastic reduction during the early stage of their infection in mosquitoes (Smith et al., 2014). Although reduced in number, *P. berghei* infection still elicits remarkable changes in *An. stephensi* metabolism as revealed by our metabolomics analyses. Glucose and trehalose are the two nutrients that are significantly reduced in mosquitoes 1 dpi. Lacking in mammalian blood (Dahlqvist, 1974), the decrease of trehalose in mosquitoes reinforces that alteration in sugar metabolism is due to the *Plasmodium* infection, not due to metabolite variation in mouse plasma ingested by mosquitoes (Li et al., 2008). The reduction of sugar might result from the reallocation of mosquito nutrients towards immune defense (Dolezal et al., 2019), or the consumption by *Plasmodium* parasite. However, our results demonstrate that the sugar reduction during *Plasmodium* infection is due to the presence of gut microbiota. Gut microbiota proliferates exponentially 24 hr after blood feeding (Pumpuni et al., 1996; Wang et al., 2012). So, the gut microbiota-mediated increase in sugar consumption might be due to the reprogramming of bacterial metabolic activity during their exponential proliferation in response to *P. berghei* infection.

Glucose is an important energy source for all living organism (Yuval et al., 1994; Zielke et al., 1984). Trehalose is the major blood sugar in insects and can be utilized after conversion to glucose (Becker et al., 1996). Our data show that increasing susceptibility to *Plasmodium* infection through glucose/trehlose supplementation before blood feeding is not due to the elevation of mosquito sugar level, suggesting that these sugars facilitate parasite infection indirectly. We then demonstrate that such elevations of parasite infection relies on the presence of microbiota and *A. bogorensis* plays an important role in this process. *Asaia* sp. is a commensal bacterium that is widespread in disease transmitting mosquitoes, including *Anopheles*, *Aedes*, and *Culex* (Crotti et al., 2009; Duguma et al., 2019; Favia et al., 2007). It mainly distributes in the midgut, salivary glands and reproductive organs of male and females and can be vertically and horizontally transmitted to the next generation (Crotti et al., 2009; Damiani et al., 2010; Favia et al., 2007). Although this bacterium is a potential target for paratransgenesis in malaria control (Shane and Grogan, 2018), we show here that *A. bogorensis* is responsible for the glucose/trehalose-mediated increasing susceptibility to *Plasmodium* infection.

Mosquito relies primarily on innate immunity to defend against invading pathogens (Cirimotich et al., 2010; Clayton et al., 2014). Here we do observe that supplementation of glucose and trehalose downregulates the expression of effectors associated with immune deficiency (Imd) pathway. However, our results reveal that suppression of Imd pathway has no influence on *P. berghei* infection, which is in agreement with the previous report (Garver et al., 2009). In addition, we find that glucose treatment changes the expression of genes encoding different domains of V-ATPase as reported (Hayek et al., 2019). Impairment of V-ATPase function facilitates *P. berghei* infection through increasing midgut pH. Moreover, we demonstrate that increase of midgut pH leads to the downregulation of Imd pathway, suggesting that midgut pH elevation is a causal factor for the suppression of immune responses. Surprisingly, we find that glucose has no effect on *P. falciparum* infection. *P. falciparum* in this study is cultured in vitro with growing conditions differing substantially from that in human blood (Carter et al., 1993; LeRoux et al., 2009). Such difference, especially increase of glucose concentration and pH value in the medium might change biological state of *P. falciparum* and makes this parasite unresponsive to glucose treatment (LeRoux et al., 2009). To simulate the in vivo infection process, we use *P. berghei* that maintained through cyclically transmitted from mice to mosquitoes in most of our study.

We next demonstrate that *A. bogorensis* participates in the regulation of midgut pH. *A. bogorensis* oxidizes acetate and lactate to carbon dioxide and water, but not ethanol to acetic acid (Yamada et al. 2000; Katsura et al. 2001; Yukphan et al. 2004; Malimas et al. 2008). Our metabolomics data show a significant decrease of lactate, citrate and succinate in the conditioned medium using glucose as carbon source, and a significant decrease of citrate, succinate and fumarate in trehalose containing medium. These results confirm that *A. bogorensis* has less capacity for acid production than other bacteria in the gut (Ano et al., 2008). Thus, it explains that expansion of *A. bogorensis* contributes to the increase of midgut pH in glucose and trehalose supplemented mosquitoes. The limitation here is that we have difficulty in detecting the metabolites changes in mosquito midguts due to the large amount of samples required by NMR. Further investigation is needed to examine the metabolite dynamics in vivo using other technology.

The pH increase triggers male gametogenesis in vitro (Billker et al., 1997; Kawamoto, 1993). We demonstrate that the pH increase in the midgut similarly promotes microgamete formation in vivo. Consistent with our finding, acidification of tsetse midgut by colonization of a commensal bacterium of *An. gambiae* inhibits trypanosome infection (Weiss et al., 2019). Gut microbiota affects parasite infection mainly through stimulating the basal immune responses or secreting anti-plasmodium metabolites (Cirimotich et al., 2011b; Romoli and Gendrin, 2018). Here we identify the third approach that gut microbiota influences *Plasmodium* infection is through modification of mosquito metabolism, thereby influencing parasite development.

Mosquito is able to differentiate different plant species and prefers to feeding on those benefiting itself most (Gouagna et al., 2014; Manda et al., 2007a; Manda et al., 2007b). Feeding on the preferred plants increases mosquito’s life span and improves its fecundity (Manda et al., 2007a; Stone et al., 2012; Yu et al., 2016). The sugars from different plant species also influence the capability of *Anopheles* to transmit malaria parasite (Ebrahimi et al., 2018; Gu et al., 2011; Hien and Dabire, 2016). Although we have no access to the natural plants in malaria endemic area, using the sugar meals simulating the composition of 4 favorite and 1 less favorite plant saps as mosquito diets, we demonstrate that the composition of sugars determines the vector capacity and such determination is associated with the ability of them to promote the proliferation of *A. bogorensis*. Notably, *P. hysterophorus* is one of the plants that mosquito feed on most frequently, but with decreased longevity and fecundity (Manda et al., 2007a; Manda et al., 2007b). Neither it promotes *P. berghei* infection compared to other mosquito preferred plants. Thus, *P. hysterophorus* might be an ideal plant species that could be expanded in malaria endemic area to reduce mosquito’s malaria transmission potential. Further study is needed to investigate the influence of natural plant saps on the composition of microbiota in field *Anopheles* mosquitoes, and examine the influence of *A.bogorensis* from field mosquitoes on malaria parasite infection outcome.

In summary, our study provides the crucial molecular insight into how interplay between glucose metabolism of *An. stephensi* and gut microbiota, *A. bogorensis*, influence *Plasmodium* infection. Our work shed light on the new role of gut microbiota in influencing mosquito metabolism and parasite infection. We also provide evidence that the different sugar composition influences mosquito’s malaria transmission potential and such influence is associated with the abundance of *A. bogorensis*. Thus, our work provides an important step toward a more comprehensive understanding of metabolic interactions between *Anopheles*, gut microbiota and malaria parasites.

## ACKNOLEDGEMENTS

We thank Dr. Jing Yuan from Xiamen University for help with gamete staining analysis in mosquito midgut. We thank Dr. Shimin Zhao and Dr. Hongyan Wang from Fudan University for comments on manuscript. We acknowledge support from the National Institutes of Health Grant (R01AI129819), National Natural Science Foundation of China (U1902211 and 31822051), the Research Fund of the State Key Laboratory of Genetic Engineering and 111 project (B13016), Fudan University to J. W.

## AUTHOR CONTRIBUTIONS

Conceptualization, M.W., and J.W., Methodology, M.W., Y.A., Y.F., S.D., G.D., H.T., and J.W.; Investigation, M.W., Y.A., S.D., Y.F; and L.G.; Formal Analysis, M.W., Y.A., H.T., and J.W., Writing Original Draft, M.W., Y.A., and J.W.; Writing, Review & Editing, M.W., Y.A., P.W., H.T., and J.W.; Visualization, M.W., Y.A., H. T., and J.W.; Funding Acquisition, J.W.; Resources, J.W., Supervision, G.D., H.T., and J.W.

## STAR METHODS

### KEY RESOURCES TABLE

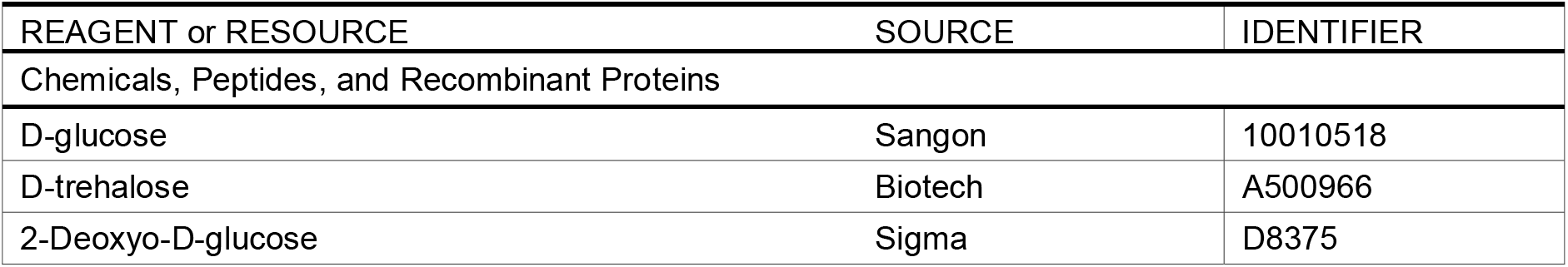
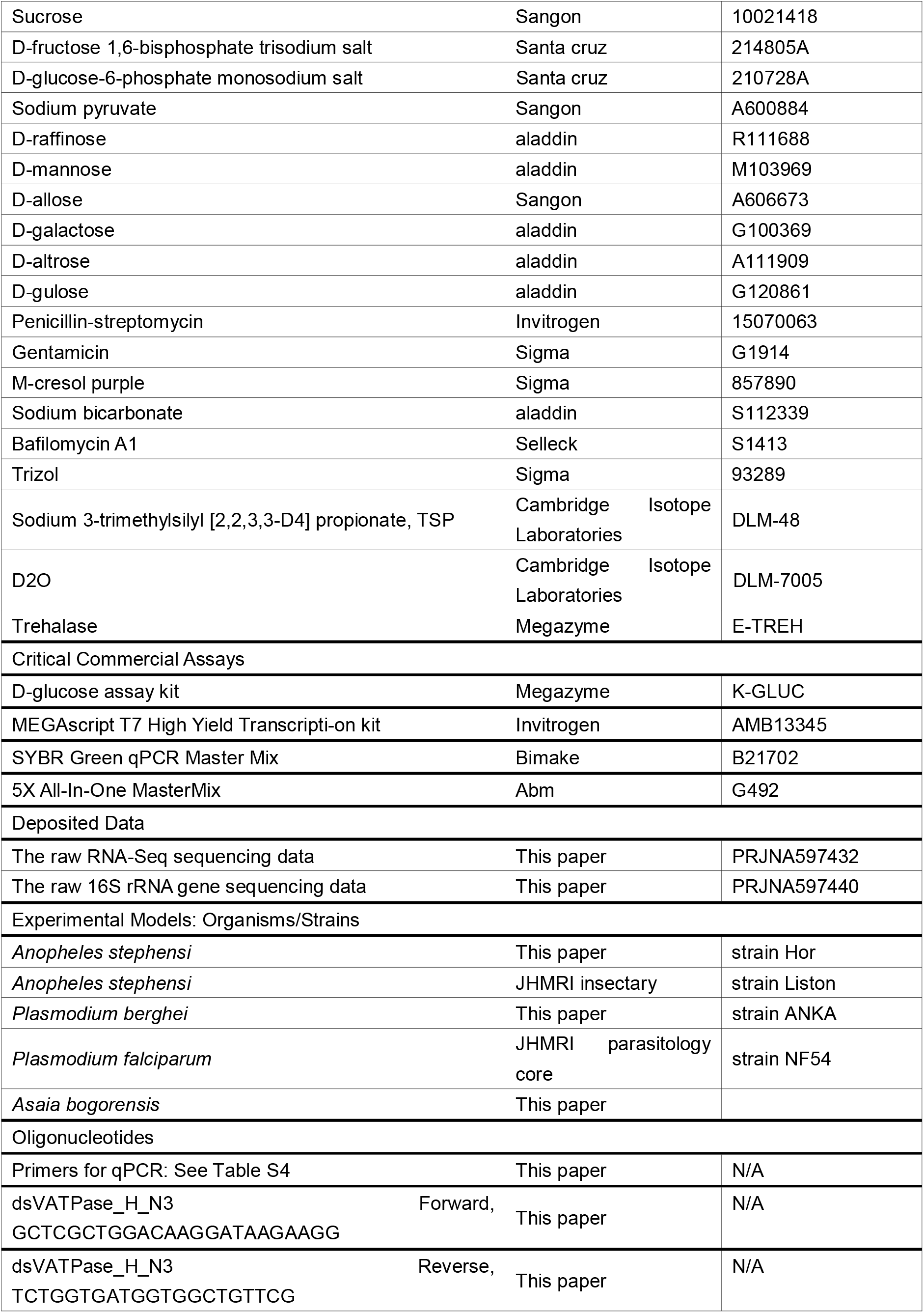
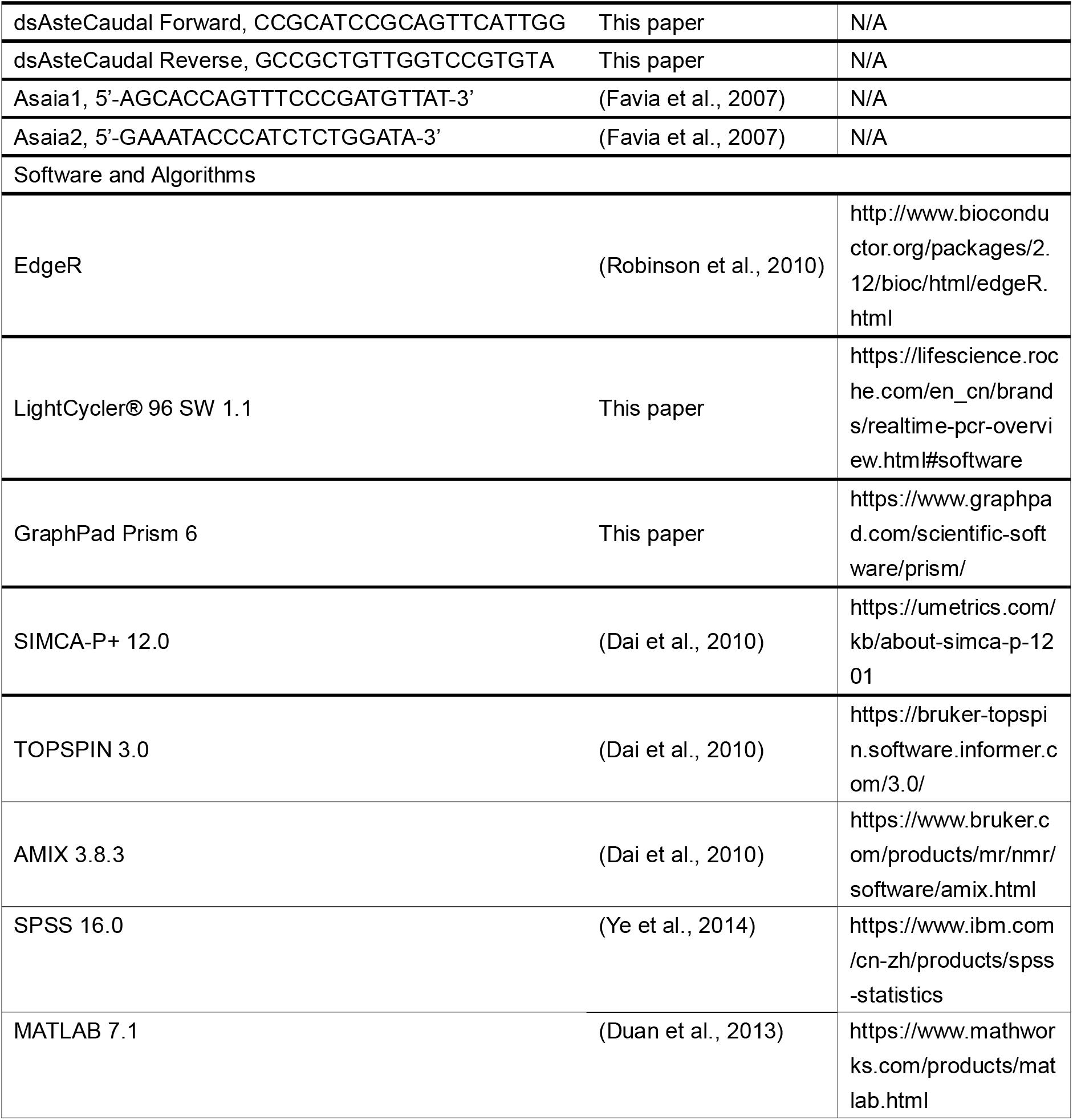

## EXPERIMENTAL MODEL and SUBJECT DETAILS

### Mosquito rearing and maintenance

*Anopheles stephensi* (strain Hor) was reared at insectary in Fudan University at 28°C, 80% relative humidity and on a 12:12 light/dark photoperiod. *Anopheles stephensi* (strain Liston) was maintained in Johns Hopkins Malaria Institute (JHMRI) insectary at 27 °C and 80% humidity with a 12 h day-night cycle. Adults were maintained on 2% (0.1 M) sucrose. Adult females were fed on anesthetized mice for a blood meal and allowed to lay eggs on wet filter paper.

### *Plasmodium berghei* maintenance

*Plasmodium berghei* ANKA parasites were maintained in BALB/c mice by serial blood passage and regular mosquito transmission (Sinden, 1997).

### *Plasmodium falciparum* culture

*P. falciparum* NF54W was provided by the Johns Hopkins Malaria Institute (JHMRI) Core Facility. It was cultured as asexual stages between 0.2 and 2% parasitemia at 37°C in 4% human erythrocytes using *P. falciparum* cultures, prepared as described (Carter et al., 1993). Parasites were kept under a gas mixture of 5% O_2_, 5%CO_2_, balanced N_2_ for up to 8 weeks according to established protocols (Trager and Jensen, 1976). Gametocytogenesis was induced by raising parasitemia > 4% and changing medium daily for 14 to 18 days obtain matured gametocyte.

### *Asaia bogorensis* culture

*Asaia bogorensis* strains were cultured in liquid medium (2% sorbitol, 0.5% peptone, 0.3% yeast extract, pH 3.5) or modified medium using 0.1 M glucose and 0.1 M trehalose as carbon source at 28°C.

## METHOD DETAILS

### *Plasmodium* infection of *An. stephensi*

Mosquitoes were blood fed on *P. berghei* infected mice when parasitemia reaches 3%-6%. Mosquitoes not fully engorged were removed 24 hr post blood meal. Fully engorged ones were maintained on 2% sucrose at 21°C until dissection. Midguts were dissected and oocysts were counted under the microscope 8 days post infection. *Anopheles stephensi* were fed on 0.02% NF54 *P. falciparum* gametocyte cultures through artificial membranes feeder at 37°C at the Johns Hopkins Malaria Institute Core Facility as described (Dong et al., 2006; Trager and Jensen, 1976). Unfed mosquitoes were removed after blood feeding, and the engorged mosquitoes were kept at 27°C for 8 days to check oocysts in the midgut. To determine oocyst numbers, midguts were dissected in PBS, stained with 0.2% mercurochrome, and oocyst numbers in midgut were counted using a phase-contrast microscope.

### Metabolite extraction

Extraction of metabolites from mosquitoes was performed as described (Chen et al., 2017a). Fifteen mosquitoes were pooled for one biological sample. Ten biological replicates from each treatment were used in the following NMR analysis. Briefly, mosquitoes were snap-frozen in liquid nitrogen and homogenized in 66% methanol solution. Supernatant was saved and pellet was re-extracted in the same methanol solution twice more. All supernatants were combined and centrifuged at 16,000 g, 4 °C for 10 min. Approximate 2 ml supernatant was transferred into a new tube. Methanol was removed under vacuum using Eppendorf Concentrator plus (Eppendorf, Germany), and the remaining liquid was lyophilized in a freeze-drier.

Extraction of metabolites from liquid medium was performed as described (Chen et al., 2017a). Briefly, 2 ml of conditioned medium was collected from the culture with *A. bogorensis* growing for 3 days. Five biological replicates from each treatment were used for NMR analysis. The collected medium was mixed with 4 ml of methanol and centrifuged at 12,000 rpm, 4°C for 10 min. Approximate 2 ml supernatant was transferred into a new tube. Methanol was removed under vacuum using Eppendorf Concentrator plus (Eppendorf, Germany), and the remaining liquid was lyophilized in a freeze-drier.

Each dried extract was reconstituted into 600 μl phosphate buffer (0.15 M, K_2_HPO_4_-NaH_2_PO_4_, pH 7.43) containing 80% D_2_O and 0.337 mM sodium 3-trimethylsilyl [2,2,3,3-D4] propionate, TSP. After vortex and centrifugation (16,099 g, 4°C, 10 min), 550 μl supernatant was transferred into a 5 mm NMR tube for NMR analysis.

### NMR spectroscopic analysis

All NMR spectra were acquired at 298 K on a Bruker Advance Ⅲ 600 MHz NMR spectrometer (600.13 MHz for proton frequency) equipped with an quaternary cryogenic inverse probe (Bruker BioSpin, Germany). ^1^D ^1^H NMR spectrum was acquired with a standard NOESYGPPR1D pulse sequence (RD−G1−90°−t1−90°−tm−G2−90°−acq) with the recycle delay (RD) of 2 s and tm of 100 ms. The 90° pulse length was adjusted to about 10 μs and 64 scans were collected into 32 k data points with spectral width of 20 ppm.

For NMR signal assignments purposes, a series of 2D NMR spectra were acquired for selected samples including ^1^H−^1^H correlation spectroscopy (COSY), ^1^H−^1^H total correlation spectroscopy (TOCSY), J-resolved spectroscopy (JRES), ^1^H−^13^C heteronuclear single quantum correlation (HSQC), and 1H−13C heteronuclear multiple bond correlation (HMBC) as previously reported (Dai et al., 2010)

Multivariate data analysis was used to analyze the metabolites from mosquito extracts 1 day post infection. Principal component analysis (PCA) and Orthogonal projection to latent structure discriminant analysis (OPLS-DA) were performed with the software package SIMCA-P+ (12.0, Umetrics, Sweden). PCA was conducted using the mean centered data, while OPLS-DA was carried out with 7-fold cross-validation using the unit-variance scaled data. Qualities of the OPLS-DA models were assessed with CV-ANOVA approach with p < 0.05 as significant level. The metabolites significantly changed were showed as back-transformed loadings plots from OPLS-DA with pearson correlation coefficients color-coded for all metabolites. In these loadings plots, the metabolites with warm color (e.g., blue) contributed more significantly to the intergroup differences than those with cold color (e.g., black).

Univariate data analysis was used to analyze the significantly changed metabolites in the conditioned medium from *A. bogorensis* culture. The relative content of metabolites in the extract of culture medium was calculated by the integral of characteristic and least-overlapping NMR signals. Comparisons between two groups were analyzed with Student’s *t* test (if criteria met) or nonparametric tests using MATLAB 7.1 software (MATLAB 7.1, Mathworks Inc., USA)(Duan et al., 2013).

### Sugar treatment

Newly emerged *An. stephensi* were fed on the different sugar diets for up to 5 days. After *Plasmodium* infection, mosquitoes were maintained on 2% sucrose unless otherwise indicated. For the assay of *P. berghei* infection in mosquito fed with different sugar diets post blood meal, newly emerged *An. stephensi* were fed on 2% sucrose until *Plasmodium* infection. Then mosquitoes were maintained on different sugar meals until oocyst number was examined. The different sugar diets are: S, 2% sucrose; G, 2% sucrose containing D-glucose with indicated concentrations; T, 2% sucrose containing 0.1 M trehalose; 2-DG, 2% sucrose containing 0.1 M D-glucose and 5 mM 2-Deoxyo-D-glucose (2-DG); HS, 10% sucrose; PYR, 2% sucrose + 0.1 M sodium pyruvate; FBP, 2% sucrose + 0.1 M D-fructose 1,6-bisphosphate trisodium salt; G6P, 2% sucrose + 0.1 M D-glucose-6-phosphate monosodium salt. Sugar diets simulating the five natural plant in West Kenya were based on the sugar composition and water content as described (Abebe, 2015; Abere and Enoghama, 2015; Fatimah and Ahmad, 2009; Manda et al., 2007a; Sowmya et al., 2019). *R.c.*, 0.4 g D-Mannose and 0.37 g D-Gulose in 100 ml water; *S.d.*, 5.91 g D-Glucose, 0.41 g D-Fructose, 1.875 g Sucrose, 1.11 g D-Mannose, 8.31 g D-Gulose, 0.55 g D-Galactose in 100 ml water; *P.h.*, 0.06 g D- Glucose, 0.26 g D-Fructose, 0.03 g Sucrose, 0.97 g D-Gulose, 0.02 g D-Allose in 100 ml water; *T.s.*, 4.21 g D- Glucose, 11.64 g D-Fructose, 0.022 g Sucrose, 0.28 g D-Mannose, 18.96 g D-Gulose, 0.28 g D-Raffinose, 1.04 g D- Altrose in 100 ml water; *L.c.*, 0.437 g D-Glucose in 100 ml water.

### Sugar measurement

Mosquito’s hemolymph and midgut sugar levels were determined as described (Kwon et al., 2015; Liu et al., 2013). Briefly, 30~40 μl hemolymph from 10 mosquitoes were collected for glucose and trehalose concentration determination. Ten midguts were pooled and homogenized in 40 μl phosphate buffered Saline (PBS) buffer. Thirty μl of midgut homogenates were collected for glucose and trehalose measurement. One third volume of individual sample was used to measure the glucose level with the Glucose Kit (Megazyme; K-GLUC). Another 1/3 volume was used to determine the trehalose level. The last 1/3 volume of the sample was used to extract and quantify the amount of genomic DNA. The concentration of glucose and trehalose were normalized to the amount of genomic DNA. Total DNA of midguts and hemolymph was extracted using Holmes Bonner method as described (Holmes and Bonner, 1973).

### Antibiotics treatment

Newly enclosed mosquitoes were administrated with fresh filtered 2% sucrose supplemented with a cocktail of antibiotics including 10 U/ml penicillin, 10 μg/ml streptomycin and 15 μg/ml gentamicin daily for up to 5 days (Dong et al., 2009). The efficacy of antibiotics treatment was examined by plating the homogenate of surface sterilized mosquitoes on LB-agars.

### Oral administration of *Asaia bogorensis*

*A. bogorensis* was isolated from the midgut of *An. stephensi* feeding on G and confirmed by 16S *rRNA* sequencing. Briefly, lysate from individual midgut was cultured in liquid medium (2% sorbitol, 0.5% peptone, 0.3% yeast extract, pH 3.5) and then plated on medium containing 0.1% D-glucose, 1.5% glycerol, 0.5% peptone, 0.5% yeast extract, 0.2% malt extract, 0.7% CaCO_3_, 1.5% agar as described (Yamada et al., 2000). To re-colonize *A. bogorensis* to the antibiotic treated *An. stephensi*, the overnight culture of this bacterium was washed twice in 0.9% NaCl and diluted to the final concentration of 10^7^~ 10^8^ cells/ml with 2% sucrose. *A. bogorensis* containing sugar solution was introduced to antibiotics treated mosquitoes and changed daily. Colonization of *A. bogorensis* was examined by qRT-PCR 2 days post treatment. Age matched normal and antibiotic treated mosquitoes were used as controls. RNA was extracted from whole mosquito or midgut using TRIzol® (Invitrogen). One μg of total RNA was used to synthesize cDNA using 5XAll-in-One MasterMix (AccuRT Genomic DNA Removal Kit, ABM, Shanghai, China). The cDNA was used as template in qPCRs with primers Asafor /Asarev. The copy number of *A. bogorensis* and *An. stephensi S7* gene was determined by standard curve. The relative abundance of *A. bogorensis* was normalized with *An. stephensi s7* gene copy number. All primers were listed in Table S4.

Quantitative real time PCR was performed by Roche LightCycler 96 Real Time PCR Detection System using the following cycling conditions: 1 cycle of 95 °C for 5 min, 40 cycles of 95 °C for 15 sec, 60 °C for 30 sec, and 72 °C for 15 sec. Melting curves (60°C to 95°C) were performed to confirm the identity of the PCR product. The data were processed and analyzed with LightCycler 96 software.

### Transcriptome analysis

Midguts of mosquitoes fed on S and G sugar meals were used for transcriptome analysis. Four mosquito midguts were pooled for one sample and three biological replicates were collected from each treatment. Total RNA were extracted using TRIzol® Reagent according the manufacturer’s instructions (Invitrogen) and sent to Majorbio (Shanghai, China) for library construction and sequencing using Illumina HiSeq xten. Clean data were aligned to the reference genome AsteS1.7 (https://www.vectorbase.org/organisms/anopheles-stephensi). The expression level of each transcript was calculated according to the fragments per kilobase of exon per million mapped reads (FRKM) method. R statistical package software EdgeR (Empirical analysis of Digital Gene Expression in R, http://www.bioconductor.org/packages/2.12/bioc/html/edgeR.html) was utilized for differential expression analysis (Robinson et al., 2010).

### Gene knockdown and quantitative reverse transcription PCR (qRT-PCR)

The cDNA clones of target genes were obtained using gene specific primers (Table S4). PCR amplicons of GFP, *vatpas*_*enh3* and *caudal* tailed with T7 promoter (TAA TAC GAC TCA CTA TAG GGA GA) were used to synthesize dsRNAs using MEGAscript T7 High Yield Transcription kit (Invitrogen). Four to six-day-old females received a total 69 nl dsRNAs (4 μg/μl) through intra-thoracically microinjection using Nanoject II microinjector (Drummond). Injected mosquitoes were allowed to recover for 4 days prior to infection (Blandin et al., 2002). Silencing efficiency was verified by qPCR 4-day post dsRNA treatment with primers listed in Table S4. The cDNA was prepared from total RNA using the 5XAll-in-One MasterMix (with AccuRT Genomic DNA Removal Kit) (ABM, China). Levels of target genes were determined by Roche LightCycler 96 Real Time PCR Detection System with SYBR Green qPCR Master Mix (Biomake). The data were processed and analyzed with LightCycler 96 software. Ribosomal gene *s7* of *An. stephensi* and *18S rRNA* of *P. berghei* were used as the internal reference (Baptista et al., 2010; Dimopoulos et al., 1996). Relative quantitation results were normalized with reference gene and analyzed by the 2^−ΔΔCt^ method (Livak and Schmittgen, 2001). Gene expression of dsRNA treated group was normalized to that of dsGFP controls.

### Gut microbiota analysis by 16S rRNA sequencing

Mosquitoes fed on S and G sugar meals were surface sterilized with 70% ethanol twice and 1× PBS twice. Total DNA of three midguts were pooled for one biological sample and extracted by the method of Holmes and Bonner as described (Holmes and Bonner, 1973). Total 5 biological replicates from each treatment were used for 16S rRNA analysis. The composition of microbiota of mosquito was analyzed using an Illumina HiSeq2500 platform in Novogene (Novogene, China) by primers targeting V3-V4 region of bacterial 16S *rRNA*. No template controls were included as controls. Following quality filtering and chimera sequences removal, a total of 772,822 reads were detected. Operational taxonomic units (OTUs) analysis were performed by Uparse software (Uparse v7.0.1001.)(Edgar, 2013). Sequences with ≥97% similarity were assigned to the same OTUs. Total 1,879 OTUs were obtained.

### Fluorescent in situ hybridization

Abdomens of females fed on S and G sugar meals were fixed and sectioned as described (Attardo et al., 2008). Slides were hybridized with *A. bogorensis* specific *16S rRNA* probe, Asaia1 (5’-AGC ACC AGT TTC CCG ATG TTA T-3’) and Asaia2 (5’-GAA ATA CCC ATC TCT GGA TA-3’) labeled with Alexa Fluor® 555 (Invitrogen). Tissues were visualized using Nikon ECLIPSE IVi microscope connected to a Nikon DIGITAL SIGHT DS-U3 digital camera.

### Midgut pH assay

Midgut pH of *An. stephensi* was determined using m-Cresol purple (Sigma). Mosquitoes were administered a sugar meal containing 2% sucrose and 0.1% m-Cresol purple (Overend et al., 2016). Midgut was dissected 6 hours post feeding and visualized immediately using a ZEISS AxioCam MRc digital camera mounted on a ZEISS AXIO Zoom.V16 stereoscope.

### Exflagellation assay

The midgut of mosquito was dissected 10 minutes after infection, and kept in PCR tube coated with heparin sodium to prevent blood clotting. After centrifugation at 2000 g for 3 min, the whole blood bolus was coated on a glass slide and stained with Giemsa. Exflagellations and exflagellation rate were calculated as described (Jones et al., 1994; Shinondo et al., 1994). Exflagellation events were counted per 10^5^ erythrocytes under a light microscope. The exflagellation rate was calculated by the ratio of the number of exflagellation events to the number of gametocytes observed per 10^5^ erythrocytes.

### Quantification and statistical Analysis

The details of statistical methods were listed in the figure legends. Data were analyzed in GraphPad Prism 6 unless indicated. The statistical analysis of metabolomics data and transcriptome data were described in the corresponding methods.

### Data and software availability

The raw RNA-Seq sequencing data were uploaded to the National Center for Biotechnology Information’s Sequence Read Archive (Accession no. PRJNA597432). The raw 16S rRNA gene sequences were uploaded to the National Center for Biotechnology Information’s Sequence Read Archive (Accession no. PRJNA597440).

## SUPPLEMENTAL INFORMATION

Supplemental information includes 4 figures and 4 tables.

## REFERENCES

Abebe, S. (2015). Fuel briquette potential of Lantana camara L. weed species and its implication for weed management and recovery of renewable energy source, in Ethiopia (Addis Ababa University).

Abere, T.A., and Enoghama, C.O. (2015). Pharmacognostic standardization and insecticidal activity of the leaves of *Tecoma stans* Juss (Bignoniaceae). Journal of Science and Practice of Pharmacy 2, 39–45.

Angrisano, F., Tan Yh Fau - Sturm, A., Sturm A Fau - McFadden, G.I., McFadden Gi Fau - Baum, J., and Baum, J. (2012). Malaria parasite colonisation of the mosquito midgut--placing the *Plasmodium* ookinete centre stage. International journal for parasitology 42(6), 519–527.

Ano, Y., Toyama, H., Adachi, O., and Matsushita, K. (2008). Energy metabolism of a unique acetic acid bacterium, *Asaia bogorensis*, that lacks ethanol oxidation activity. Bioscience, biotechnology, and biochemistry 72, 989–997.

Attardo, G.M., Lohs, C., Heddi, A., Alam, U.H., Yildirim, S., and Aksoy, S. (2008). Analysis of milk gland structure and function in *Glossina morsitans*: milk protein production, symbiont populations and fecundity. Journal of insect physiology 54, 1236–1242.

Bahia, A.C., Dong, Y., Blumberg, B.J., Mlambo, G., Tripathi, A., BenMarzouk Hidalgo, O.J., Chandra, R., and Dimopoulos, G. (2014). Exploring *Anopheles* gut bacteria for *Plasmodium* blocking activity. Environmental microbiology 16, 2980–2994.

Baptista, F.G., Pamplona, A., Pena, A.C., Mota, M.M., Pied, S., and Vigário, A.M. (2010). Accumulation of *Plasmodium berghei*-infected red blood cells in the brain is crucial for the development of cerebral malaria in mice. Infection and immunity 78, 4033–4039.

Becker, A., Schlöder, P., Steele, J., and Wegener, G. (1996). The regulation of trehalose metabolism in insects. Experientia 52, 433–439.

Bennink, S., Kiesow, M.J., and Pradel, G. (2016). The development of malaria parasites in the mosquito midgut. Cellular microbiology 18, 905–918.

Billker, O., Dechamps, S., Tewari, R., Wenig, G., Franke-Fayard, B., and Brinkmann, V. (2004). Calcium and a calcium-dependent protein kinase regulate gamete formation and mosquito transmission in a malaria parasite. Cell 117, 503–514.

Billker, O., Shaw, M.K., Margos, G., and Sinden, R.E. (1997). The roles of temperature, pH and mosquito factors as triggers of male and female gametogenesis of *Plasmodium berghei* in vitro. Parasitology 115, 1–7.

Blandin, S., Moita, L.F., Köcher, T., Wilm, M., Kafatos, F.C., and Levashina, E.A. (2002). Reverse genetics in the mosquito *Anopheles gambiae*: targeted disruption of the Defensin gene. EMBO reports 3, 852–856.

Boudko, D.Y., Moroz, L.L., Harvey, W.R., and Linser, P.J. (2001a). Alkalinization by chloride/bicarbonate pathway in larval mosquito midgut. Proceedings of the National Academy of Sciences 98, 15354.

Boudko, D.Y., Moroz, L.L., Linser, P.J., Trimarchi, J.R., Smith, P.J., and Harvey, W.R. (2001b). In situ analysis of pH gradients in mosquito larvae using non-invasive, self-referencing, pH-sensitive microelectrodes. Journal of Experimental Biology 204, 691–699.

Carter, R., and Nijhout, M. (1977). Control of gamete formation (exflagellation) in malaria parasites. Science 195, 407–409.

Carter, R., Ranford-Cartwright, L., and Alano, P. (1993). The culture and preparation of gametocytes of *Plasmodium falciparum* for immunochemical, molecular, and mosquito infectivity studies. Methods in molecular biology 21, 67–88.

Chen, F., Liu, C., Zhang, J., Lei, H., Li, H.P., Liao, Y.C., and Tang, H. (2017a). Combined Metabonomic and Quantitative RT-PCR Analyses Revealed Metabolic Reprogramming Associated with *Fusarium graminearum* Resistance in Transgenic *Arabidopsis thaliana*. Frontiers in plant science 8, 2177.

Chen, S., Blom, J., and Walker, E.D. (2017b). Genomic, physiologic, and symbiotic characterization of *Serratia marcescens* strains Isolated from the mosquito *Anopheles stephensi*. Front Microbiol 8, 1483.

Cirimotich, C.M., Dong, Y., Clayton, A.M., Sandiford, S.L., Souza-Neto, J.A., Mulenga, M., and Dimopoulos, G. (2011a). Natural microbe-mediated refractoriness to *Plasmodium* infection in *Anopheles gambiae*. Science 332, 855–858.

Cirimotich, C.M., Dong, Y., Garver, L.S., Sim, S., and Dimopoulos, G. (2010). Mosquito immune defenses against Plasmodium infection. Developmental & Comparative Immunology 34, 387–395.

Cirimotich, C.M., Ramirez, J.L., and Dimopoulos, G. (2011b). Native microbiota shape insect vector competence for human pathogens. Cell host & microbe 10, 307–310.

Clayton, A.M., Cirimotich, C.M., Dong, Y., and Dimopoulos, G. (2013). Caudal is a negative regulator of the *Anopheles* IMD pathway that controls resistance to *Plasmodium falciparum* infection. Developmental and comparative immunology 39, 323–332.

Clayton, A.M., Dong, Y., and Dimopoulos, G. (2014). The *Anopheles* innate immune system in the defense against malaria infection. J Innate Immun 6, 169–181.

Corby-Harris, V., Drexler, A., De Jong, L.W., Antonova, Y., Pakpour, N., Ziegler, R., Ramberg, F., Lewis, E.E., Brown, J.M., and Luckhart, S. (2010). Activation of Akt signaling reduces the prevalence and intensity of malaria parasite infection and lifespan in *Anopheles stephensi* mosquitoes. PLoS pathogens 6, e1001003.

Crotti, E., Damiani, C., Pajoro, M., Gonella, E., Rizzi, A., Ricci, I., Negri, I., Scuppa, P., Rossi, P., Ballarini, P., et al. (2009). *Asaia*, a versatile acetic acid bacterial symbiont, capable of cross-colonizing insects of phylogenetically distant genera and orders. Environmental microbiology 11, 3252–3264.

Dahlqvist, A. (1974). Enzyme deficiency and malabsorption of carbohydrates. Sugars in Nutrition HL Sipple & KW McNutt, eds.

Dai, H., Xiao, C., Liu, H., and Tang, H. (2010). Combined NMR and LC-MS analysis reveals the metabonomic changes in *Salvia miltiorrhiza* Bunge induced by water depletion. Journal of proteome research 9, 1460–1475.

Damiani, C., Ricci, I., Crotti, E., Rossi, P., Rizzi, A., Scuppa, P., Capone, A., Ulissi, U., Epis, S., and Genchi, M. (2010). Mosquito-bacteria symbiosis: the case of *Anopheles gambiae* and *Asaia*. Microbial ecology 60, 644–654.

Deligianni, E., Morgan, R.N., Bertuccini, L., Kooij, T.W., Laforge, A., Nahar, C., Poulakakis, N., Schuler, H., Louis, C., Matuschewski, K., et al. (2011). Critical role for a stage-specific actin in male exflagellation of the malaria parasite. Cellular microbiology *13*, 1714–1730.

Dimopoulos, G., Richman, A., Della Torre, A., Kafatos, F.C., and Louis, C. (1996). Identification and characterization of differentially expressed cDNAs of the vector mosquito, *Anopheles gambiae*. Proceedings of the National Academy of Sciences 93, 13066–13071.

Dolezal, T., Krejcova, G., Bajgar, A., Nedbalova, P., and Strasser, P. (2019). Molecular regulations of metabolism during immune response in insects. Insect biochemistry and molecular biology 109, 31–42.

Domingos, A., Pinheiro-Silva, R., Couto, J., do Rosario, V., and de la Fuente, J. (2017). The *Anopheles gambiae* transcriptome - a turning point for malaria control. Insect molecular biology 26, 140–151.

Dong, Y., Aguilar, R., Xi, Z., Warr, E., Mongin, E., and Dimopoulos, G. (2006). *Anopheles gambiae* immune responses to human and rodent *Plasmodium parasite* species. PLoS pathogens 2, e52.

Dong, Y., Manfredini, F., and Dimopoulos, G. (2009). Implication of the mosquito midgut microbiota in the defense against malaria parasites. PLoS pathogens 5, e1000423.

Duan, Y., An, Y., Li, N., Liu, B., Wang, Y., and Tang, H. (2013). Multiple univariate data analysis reveals the inulin effects on the high-fat-diet induced metabolic alterations in rat myocardium and testicles in the preobesity state. Journal of proteome research 12, 3480–3495.

Duguma, D., Hall, M.W., Smartt, C.T., Debboun, M., and Neufeld, J.D. (2019). Microbiota variations in *Culex nigripalpus* disease vector mosquito of West Nile virus and Saint Louis Encephalitis from different geographic origins. PeerJ 6, e6168.

Ebrahimi, B., Jackson, B.T., Guseman, J.L., Przybylowicz, C.M., Stone, C.M., and Foster, W.A. (2018). Alteration of plant species assemblages can decrease the transmission potential of malaria mosquitoes. The Journal of applied ecology 55, 841–851.

Edgar, R.C. (2013). UPARSE: highly accurate OTU sequences from microbial amplicon reads. Nat Methods 10, 996–998.

Fatimah, H., and Ahmad, T. (2009). Phenology of *Parthenium hysterophorus*–a key factor for the success of its invasion. Adv Environ Biol 3, 150–156.

Favia, G., Ricci, I., Damiani, C., Raddadi, N., Crotti, E., Marzorati, M., Rizzi, A., Urso, R., Brusetti, L., Borin, S., et al. (2007). Bacteria of the genus *Asaia* stably associate with *Anopheles stephensi,* an Asian malarial mosquito vector. Proceedings of the National Academy of Sciences of the United States of America 104, 9047–9051.

Foster, W.A. (1995). Mosquito sugar feeding and reproductive energetics. Annual review of entomology 40, 443–474.

Gaio, A.d.O., Gusmão, D.S., Santos, A.V., Berbert-Molina, M.A., Pimenta, P.F.P., and Lemos, F.J.A. (2011). Contribution of midgut bacteria to blood digestion and egg production in *Aedes aegypti* (diptera: culicidae) (L.). Parasites & vectors 4, 105.

Gardner, M.J., Hall, N., Fung, E., White, O., Berriman, M., Hyman, R.W., Carlton, J.M., Pain, A., Nelson, K.E., Bowman, S., et al. (2002). Genome sequence of the human malaria parasite *Plasmodium falciparum*. Nature 419, 498–511.

Garver, L.S., Dong, Y., and Dimopoulos, G. (2009). Caspar controls resistance to *Plasmodium falciparum* in diverse anopheline species. PLoS pathogens 5, e1000335.

Gonzalez-Ceron, L., Santillan, F., Rodriguez, M.H., Mendez, D., and Hernandez-Avila, J.E. (2003). Bacteria in midguts of field-collected *Anopheles albimanus* block *Plasmodium vivax* sporogonic development. Journal of medical entomology 40, 371–374.

Gouagna, L.C., Kerampran, R., Lebon, C., Brengues, C., Toty, C., Wilkinson, D.A., Boyer, S., and Fontenille, D. (2014). Sugar-source preference, sugar intake and relative nutritional benefits in *Anopheles arabiensis* males. Acta tropica 132, S70–79.

Gu, W., Müller, G., Schlein, Y., Novak, R.J., and Beier, J.C. (2011). Natural plant sugar sources of *Anopheles* mosquitoes strongly impact malaria transmission potential. PloS one 6, e15996.

Guttery, D.S., Poulin, B., Ramaprasad, A., Wall, R.J., Ferguson, D.J., Brady, D., Patzewitz, E.M., Whipple, S., Straschil, U., Wright, M.H., et al. (2014). Genome-wide functional analysis of *Plasmodium* protein phosphatases reveals key regulators of parasite development and differentiation. Cell host & microbe 16, 128–140.

Hayek, S.R., Rane, H.S., and Parra, K.J. (2019). Reciprocal regulation of V-ATPase and glycolytic pathway elements in health and disease. Frontiers in physiology 10, 127.

Hien, D.F., and Dabire, K.R. (2016). Plant-mediated effects on mosquito capacity to transmit human malaria. PLoS pathogens 12, e1005773.

Holmes, D.S., and Bonner, J. (1973). Preparation, molecular weight, base composition, and secondary structure of giant nuclear ribonucleic acid. Biochemistry 12, 2330–2338.

Impoinvil, D.E., Kongere, J.O., Foster, W.A., Njiru, B.N., Killeen, G.F., Githure, J.I., Beier, J.C., Hassanali, A., and Knols, B.G. (2004). Feeding and survival of the malaria vector *Anopheles gambiae* on plants growing in Kenya. Medical and veterinary entomology 18, 108–115.

Itani, S., Torii, M., and Ishino, T. (2014). D-Glucose concentration is the key factor facilitating liver stage maturation of *Plasmodium*. Parasitology international 63, 584–590.

Jagadeshwaran, U., Onken, H., Hardy, M., Moffett, S.B., and Moffett, D.F. (2010). Cellular mechanisms of acid secretion in the posterior midgut of the larval mosquito (*Aedes aegypti*). The Journal of experimental biology 213, 295–300.

Joët, T., Eckstein-Ludwig, U., Morin, C., and Krishna, S. (2003). Validation of the hexose transporter of *Plasmodium falciparum* as a novel drug target. Proceedings of the National Academy of Sciences 100, 7476–7479.

Jones, I.W., Denholm, A.A., Ley, S.V., Lovell, H., Wood, A., and Sinden, R.E. (1994). Sexual development of malaria parasites is inhibited in vitro by the Neem extract Azadirachtin, and its semi-synthetic analogues. FEMS microbiology letters 120, 267–273.

Kawamoto, F. (1993). Ionic regulation and signal transduction system involved in the induction of gametogenesis in malaria parasites. Annals of the New York Academy of Sciences 707, 431–434.

Kawamoto, F., Alejo-Blanco, R., Fleck, S.L., and Sinden, R.E. (1991). *Plasmodium berghei*: ionic regulation and the induction of gametogenesis. Experimental parasitology 72, 33–42.

Kirk, K. (2001). Membrane transport in the malaria-infected erythrocyte. Physiol Rev 81, 495–537.

Kwon, Y., Song, W., Droujinine, I.A., Hu, Y., Asara, J.M., and Perrimon, N. (2015). Systemic organ wasting induced by localized expression of the secreted insulin/IGF antagonist ImpL2. Developmental cell 33, 36–46.

Lanfrancotti, A., Bertuccini, L., Silvestrini, F., and Alano, P. (2007). *Plasmodium falciparum*: mRNA co-expression and protein co-localisation of two gene products upregulated in early gametocytes. Experimental parasitology 116, 497–503.

LeRoux, M., Lakshmanan, V., and Daily, J.P. (2009). *Plasmodium falciparum* biology: analysis of in vitro versus in vivo growth conditions. Trends in parasitology 25, 474–481.

Li, J.V., Wang, Y., Saric, J., Nicholson, J.K., Dirnhofer, S., Singer, B.H., Tanner, M., Wittlin, S., Holmes, E., and Utzinger, J. (2008). Global metabolic responses of NMRI mice to an experimental *Plasmodium berghei* infection. Journal of proteome research 7, 3948–3956.

Linser, P.J., Smith, K.E., Seron, T.J., and Neira Oviedo, M. (2009). Carbonic anhydrases and anion transport in mosquito midgut pH regulation. The Journal of experimental biology 212, 1662–1671.

Liu, K., Dong, Y., Huang, Y., Rasgon, J.L., and Agre, P. (2013). Impact of trehalose transporter knockdown on *Anopheles gambiae* stress adaptation and susceptibility to *Plasmodium falciparum* infection. Proceedings of the National Academy of Sciences of the United States of America 110, 17504–17509.

Livak, K.J., and Schmittgen, T.D. (2001). Analysis of relative gene expression data using real-time quantitative PCR and the 2^−ΔΔ^CT method. methods 25, 402–408.

Luckhart, S., Giulivi, C., Drexler, A.L., Antonovakoch, Y., Sakaguchi, D., Napoli, E., Wong, S., Price, M.S., Eigenheer, R., and Phinney, B.S. (2013). Sustained activation of Akt elicits mitochondrial dysfunction to block *Plasmodium falciparum* Infection in the mosquito host. PLoS pathogens 9, e1003180–e1003180.

Magnarelli, L.A. (1978). Nectar-feeding by female mosquitoes and its relation to follicular development and parity. Journal of medical entomology 14, 527–530.

Mancio-Silva, L., Slavic, K., Ruivo, M.T.G., Grosso, A.R., Modrzynska, K.K., Vera, I.M., Sales-Dias, J., Gomes, A.R., MacPherson, C.R., and Crozet, P. (2017). Nutrient sensing modulates malaria parasite virulence. Nature 547, 213.

Manda, H., Gouagna, L.C., Foster, W.A., Jackson, R.R., Beier, J.C., Githure, J.I., and Hassanali, A. (2007a). Effect of discriminative plant-sugar feeding on the survival and fecundity of Anopheles gambiae. Malar J 6, 113.

Manda, H., Gouagna, L.C., Nyandat, E., Kabiru, E.W., Jackson, R.R., Foster, W.A., Githure, J.I., Beier, J.C., and Hhassanali, A. (2007b). Discriminative feeding behaviour of *Anopheles gambiae* s.s. on endemic plants in western Kenya. Medical and veterinary entomology 21, 103–111.

Marques, S.R., Ramakrishnan, C., Carzaniga, R., Blagborough, A.M., Delves, M.J., Talman, A.M., and Sinden, R.E. (2015). An essential role of the basal body protein SAS-6 in *Plasmodium* male gamete development and malaria transmission. Cellular microbiology 17, 191–206.

McKee, R.W., Ormsbee, R.A., Anfinsen, C.B., Geiman, Q.M., and Ball, E.G. (1946). Studies on malarial parasites: VI. The chemistry and metabolism of normal and parasitized (*P. knowlesi*) monkey blood. The Journal of experimental medicine 84, 569.

Meireles, P., Sales-Dias, J., Andrade, C.M., Mello-Vieira, J., Mancio-Silva, L., Simas, J.P., Staines, H.M., and Prudêncio, M. (2016). GLUT1-mediated glucose uptake plays a crucial role during *Plasmodium* hepatic infection. Cellular microbiology 19, e12646.

Minard, G., Tran, F.H., Raharimalala, F.N., Hellard, E., Ravelonandro, P., Mavingui, P., and Valiente Moro, C. (2013). Prevalence, genomic and metabolic profiles of *Acinetobacter* and *Asaia* associated with field-caught *Aedes albopictus* from Madagascar. FEMS microbiology ecology 83, 63–73.

Nayar, J., and Sauerman Jr, D. (1971). Physiological effects of carbohydrates on survival, metabolism, and flight potential of female *Aedes taeniorhynchus*. Journal of insect physiology 17, 2221–2233.

Nayar, J.K., and Sauerman, D.M., Jr. (1975). The effects of nutrition on survival and fecundity in Florida mosquitoes. Part 3. Utilization of blood and sugar for fecundity. Journal of medical entomology *12*, 220–225.

Olivieri, A., Bertuccini, L., Deligianni, E., Franke-Fayard, B., Curra, C., Siden-Kiamos, I., Hanssen, E., Grasso, F., Superti, F., Pace, T., et al. (2015). Distinct properties of the egress-related osmiophilic bodies in male and female gametocytes of the rodent malaria parasite *Plasmodium berghei*. Cellular microbiology 17, 355–368.

Olszewski, K.L., and Llinas, M. (2011). Central carbon metabolism of *Plasmodium* parasites. Molecular and biochemical parasitology 175, 95–103.

Overend, G., Luo, Y., Henderson, L., Douglas, A.E., Davies, S.A., and Dow, J.A. (2016). Molecular mechanism and functional significance of acid generation in the *Drosophila* midgut. Sci Rep 6, 27242.

Patrick, M.L., Aimanova, K., Sanders, H.R., and Gill, S.S. (2006). P-type Na+/K+-ATPase and V-type H+-ATPase expression patterns in the osmoregulatory organs of larval and adult mosquito *Aedes aegypti*. The Journal of experimental biology 209, 4638–4651.

Pietri, J.E., Pakpour, N., Napoli, E., Song, G., Pietri, E., Potts, R., Cheung, K.W., Walker, G., Riehle, M.A., and Starcevich, H. (2016). Two insulin-like peptides differentially regulate malaria parasite infection in the mosquito through effects on intermediary metabolism. Biochemical Journal 473, 3487–3503.

Ponzi, M., Siden-Kiamos, I., Bertuccini, L., Curra, C., Kroeze, H., Camarda, G., Pace, T., Franke-Fayard, B., Laurentino, E.C., Louis, C., et al. (2009). Egress of *Plasmodium berghei* gametes from their host erythrocyte is mediated by the MDV-1/PEG3 protein. Cellular microbiology 11, 1272–1288.

Pumpuni, C.B., Demaio, J., Kent, M., Davis, J.R., and Beier, J.C. (1996). Bacterial population dynamics in three anopheline species: the impact on *Plasmodium* sporogonic development. The American journal of tropical medicine and hygiene 54, 214–218.

Ricci, I., Damiani, C., Capone, A., DeFreece, C., Rossi, P., and Favia, G. (2012). Mosquito/microbiota interactions: from complex relationships to biotechnological perspectives. Current opinion in microbiology 15, 278–284.

Robinson, M.D., McCarthy, D.J., and Smyth, G.K. (2010). edgeR: a Bioconductor package for differential expression analysis of digital gene expression data. BIOINFORMATICS 26, 139–140.

Romoli, O., and Gendrin, M. (2018). The tripartite interactions between the mosquito, its microbiota and *Plasmodium*. Parasites & vectors 11, 200.

Roth, E.F., Jr. (1987). Malarial parasite hexokinase and hexokinase-dependent glutathione reduction in the *Plasmodium falciparum*-infected human erythrocyte. The Journal of biological chemistry 262, 15678–15682.

Saliba, K.J., Krishna, S., and Kirk, K. (2004). Inhibition of hexose transport and abrogation of pH homeostasis in the intraerythrocytic malaria parasite by an O-3-hexose derivative. FEBS letters 570, 93–96.

Samaddar, N., Paul, A., Chakravorty, S., Chakraborty, W., Mukherjee, J., Chowdhuri, D., and Gachhui, R. (2011). Nitrogen fixation in *Asaia* sp. (family Acetobacteraceae). Current microbiology 63, 226–231.

Sebastian, S., Brochet, M., Collins, M.O., Schwach, F., Jones, M.L., Goulding, D., Rayner, J.C., Choudhary, J.S., and Billker, O. (2012). A *Plasmodium* calcium-dependent protein kinase controls zygote development and transmission by translationally activating repressed mRNAs. Cell host & microbe 12, 9–19.

Shane, J.L., and Grogan, C.L. (2018). Blood meal-induced inhibition of vector-borne disease by transgenic microbiota. Nature communications 9, 4127.

Shinondo, C., Lanners, H., Lowrie, R., and Wiser, M. (1994). Effect of pyrimethamine resistance on sporogony in a *Plasmodium berghei*/*Anopheles stephensi* model. Experimental parasitology 78, 194–202.

Sinden, R.E. (1997). Infection of mosquitoes with rodent malaria. In The molecular biology of insect disease vectors (Springer), pp. 67–91.

Slavic, K., Delves, M.J., Prudencio, M., Talman, A.M., Straschil, U., Derbyshire, E.T., Xu, Z., Sinden, R.E., Mota, M.M., Morin, C., et al. (2011). Use of a selective inhibitor to define the chemotherapeutic potential of the plasmodial hexose transporter in different stages of the parasite’s life cycle. Antimicrobial agents and chemotherapy *55*, 2824–2830.

Smith, R.C., Vega-Rodriguez, J., and Jacobs-Lorena, M. (2014). The *Plasmodium* bottleneck: malaria parasite losses in the mosquito vector. Memorias do Instituto Oswaldo Cruz 109, 644–661.

Song, X., Wang, M., Dong, L., Zhu, H., and Wang, J. (2018). PGRP-LD mediates *A. stephensi* vector competency by regulating homeostasis of microbiota-induced peritrophic matrix synthesis. PLoS pathogens *14*, e1006899.

Sowmya, P., Vanaja, M., Sunita, V., and Raghuram Reddy, P. (2019). Genetic advance for physiological parameters in two high yielding castor (*Ricinus communis* L.) genotypes under irrigation and moisture stress. Journal of Pharmacognosy and Phytochemistry *SP2*, 179–184.

Stanway, R.R., Bushell, E., Chiappino-Pepe, A., Roques, M., Sanderson, T., Franke-Fayard, B., Caldelari, R., Golomingi, M., Nyonda, M., and Pandey, V. (2019). Genome-scale identification of essential metabolic processes for targeting the *Plasmodium* liver stage. Cell 179, 1112–1128.e1126.

Stone, C.M., Jackson, B.T., and Foster, W.A. (2012). Effects of plant-community composition on the vectorial capacity and fitness of the malaria mosquito *Anopheles gambiae*. The American journal of tropical medicine and hygiene 87, 727–736.

Sturm, A., Mollard, V., Cozijnsen, A., Goodman, C.D., and McFadden, G.I. (2015). Mitochondrial ATP synthase is dispensable in blood-stage *Plasmodium berghei* rodent malaria but essential in the mosquito phase. Proceedings of the National Academy of Sciences of the United States of America 112, 10216–10223.

Talman, A.M., Lacroix, C., Marques, S.R., Blagborough, A.M., Carzaniga, R., Menard, R., and Sinden, R.E. (2011). PbGEST mediates malaria transmission to both mosquito and vertebrate host. Molecular microbiology 82, 462–474.

Theriot, C.M., Koenigsknecht, M.J., Carlson, P.E., Jr., Hatton, G.E., Nelson, A.M., Li, B., Huffnagle, G.B., J, Z.L., and Young, V.B. (2014). Antibiotic-induced shifts in the mouse gut microbiome and metabolome increase susceptibility to *Clostridium difficile* infection. Nature communications 5, 3114.

Trager, W., and Jensen, J.B. (1976). Human malaria parasites in continuous culture. Science 193, 673–675.

Wang, A., Huen, S.C., Luan, H.H., Baker, K., Rinder, H., Booth, C.J., and Medzhitov, R. (2018). Glucose metabolism mediates disease tolerance in cerebral malaria. Proceedings of the National Academy of Sciences of the United States of America 115, 11042–11047.

Wang, S., Ghosh, A.K., Bongio, N., Stebbings, K.A., Lampe, D.J., and Jacobs-Lorena, M. (2012). Fighting malaria with engineered symbiotic bacteria from vector mosquitoes. Proceedings of the National Academy of Sciences of the United States of America 109, 12734–12739.

Weiss, B.L., Maltz, M.A., Vigneron, A., Wu, Y., Walter, K.S., O’Neill, M.B., Wang, J., and Aksoy, S. (2019). Colonization of the tsetse fly midgut with commensal *Kosakonia cowanii* Zambiae inhibits trypanosome infection establishment. PLoS pathogens 15, e1007470.

Yamada, Y., Katsura, K., Kawasaki, H., Widyastuti, Y., Saono, S., Seki, T., Uchimura, T., and Komagata, K. (2000). *Asaia bogorensis* gen. nov., sp. nov., an unusual acetic acid bacterium in the alpha*-Proteobacteria*. International journal of systematic and evolutionary microbiology 50 Pt 2, 823–829.

Ye, Y., An, Y., Li, R., Mu, C., and Wang, C. (2014). Strategy of metabolic phenotype modulation in Portunus trituberculatus exposed to low salinity. Journal of agricultural and food chemistry 62, 3496–3503.

Yeoh, L.M., Goodman, C.D., Mollard, V., McFadden, G.I., and Ralph, S.A. (2017). Comparative transcriptomics of female and male gametocytes in *Plasmodium berghei* and the evolution of sex in alveolates. BMC Genomics 18, 734.

Yoshimori, T., Yamamoto, A., Moriyama, Y., Futai, M., and Tashiro, Y. (1991). Bafilomycin A1, a specific inhibitor of vacuolar-type H (+)-ATPase, inhibits acidification and protein degradation in lysosomes of cultured cells. Journal of Biological Chemistry 266, 17707–17712.

Yu, B.T., Ding, Y.M., Mo, X.C., Liu, N., Li, H.J., and Mo, J.C. (2016). Survivorship and fecundity of *Culex pipiens* pallens feeding on flowering plants and seed pods with differential preferences. Acta tropica 155, 51–57.

Yuda, M., Iwanaga, S., Kaneko, I., and Kato, T. (2015). Global transcriptional repression: An initial and essential step for *Plasmodium* sexual development. Proceedings of the National Academy of Sciences of the United States of America 112, 12824–12829.

Yuval, B., Holliday-Hanson, M.L., and Washing, R.K. (1994). Energy budget of swarming male mosquitoes. Ecological Entomology 19, 74–78.

Zhuang, Z., Linser, P.J., and Harvey, W.R. (1999). Antibody to H(+) V-ATPase subunit E colocalizes with portasomes in alkaline larval midgut of a freshwater mosquito (*Aedes aegypti*). The Journal of experimental biology 202, 2449–2460.

Zielke, H.R., Zielke, C.L., and Ozand, P.T. (1984). Glutamine: a major energy source for cultured mammalian cells. Paper presented at: Federation proceedings.

