## Supplemental Figure 1-4 for "Glucose-mediated expansion of a gut commensal bacterium promotes *Plasmodium* infection through alkalizing mosquito midgut"

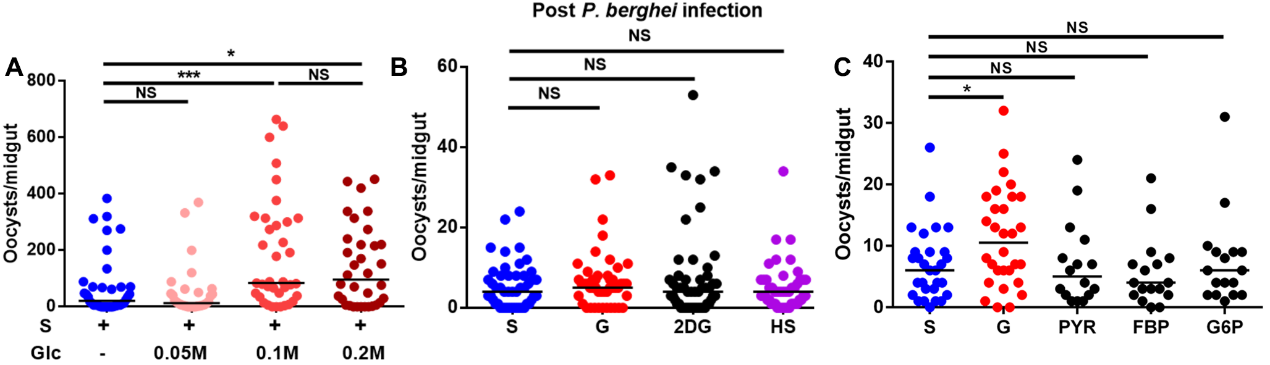

**Figure S1. The influence of sugar treatment on vector compentence of *An. stephensi*. Related to figure 1.**

(A) *P. berghei* oocyst intensity in mosquito fed with 2% sucrose containing 0 M, 0.05 M, 0.1 M and 0.2 M glucose, respectively.

(B) *P. berghei* oocyst intensity in mosquito fed with S, G, 2-DG and HS diets post blood meal, respectively.

(C) *P. berghei* oocyst intensity in mosquito fed with S, G, PYR, FBP and G6P diets, respectively.

Each dot represents an individual mosquito and horizontal lines represent the medians. Results shown in A-C were pooled from at least 2 independent experiments. Significance was determined by Mann-Whitney test. NS, not significant, *p <0.05, and ***p<0.001. S, 2% sucrose; G, 2% sucrose+0.1 M glucose; T, 2% sucrose+0.1 M trehalose; 2-DG, 2% sucrose+0.1 M glucose+5 mM 2-DG; HS, 10% sucrose; PYR, 2% sucrose+0.1 M sodium pyruvate; FBP, 2% sucrose+0.1 M D-fructose 1,6-bisphosphate trisodium; G6P, 2% sucrose+0.1 M D-glucose-6-phosphate monosodium.

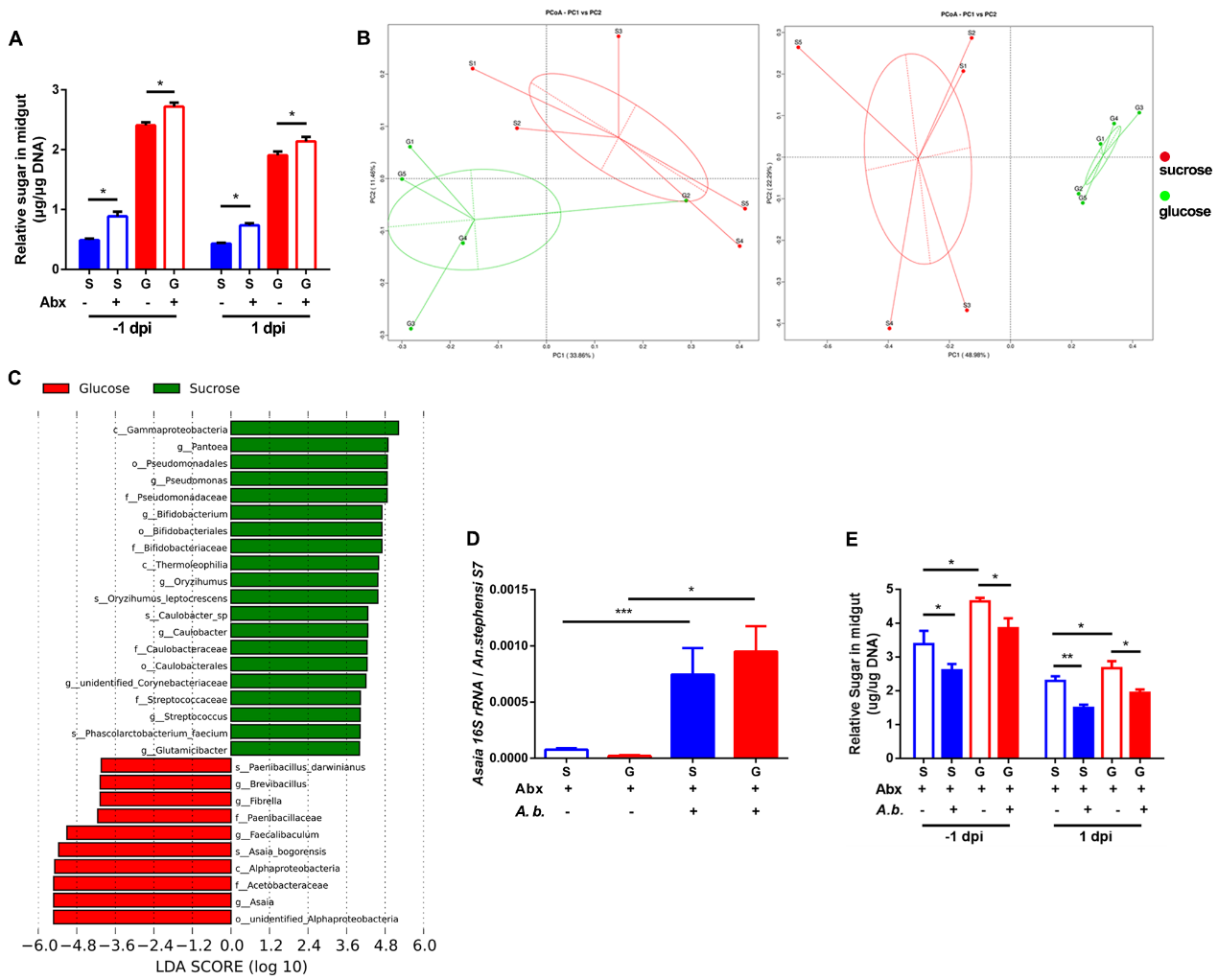
**Figure S2. The influence of *Asaia bogorensis* on sugar consumption and *P. berghei* infection. Related to figure 3.**

(A) The relative concentration of total sugar (glucose+trehalose) in the midgut of antibiotics treated mosquitoes fed on different sugar diets 1 day prior to (-1 dpi) and 1 day (1 dpi) post infection, respectively.

(B) Component Analysis (PCA) of bacterial composition by unweighted (left panel) and weighted (right panel) unifrac analyses. Each plot represents 5 midguts as a biological repeat.

(C) LEfSe analysis of mosquito midgut microbiota fed on S and G. LDA scores showed significant bacterial differences within groups at the different levels.

Green, mosquito fed on 2% sucorse; Red, mosquito fed on 2% sucrose+ 0.1 M glucose.

(D) The abundance of *A. bogorensis* in the midgut of *An. stephensi* determined by qPCR.

(E) The relative concentration of total sugar (glucose and trehalose) in the midgut of mosquitoes1 day prior to (-1 dpi) and 1 day (1 dpi) post infection, respectively.

Glucose and trehalose concentrations in A and E were normalized to genomic DNA extracted from midguts. Significance in A, D and E was determined by Student’s *t* test. Data were shown as mean ± SEM (A and E, *n*=5, D, *n*=10). NS, no significance, *, p <0.05, **, p<0.01and ***, p<0.001S, 2% sucrose; G, 2% sucrose+0.1 M glucose, Abx, antibiotics treatment, *A. b.*, *A. bogorensis* re-colonization.

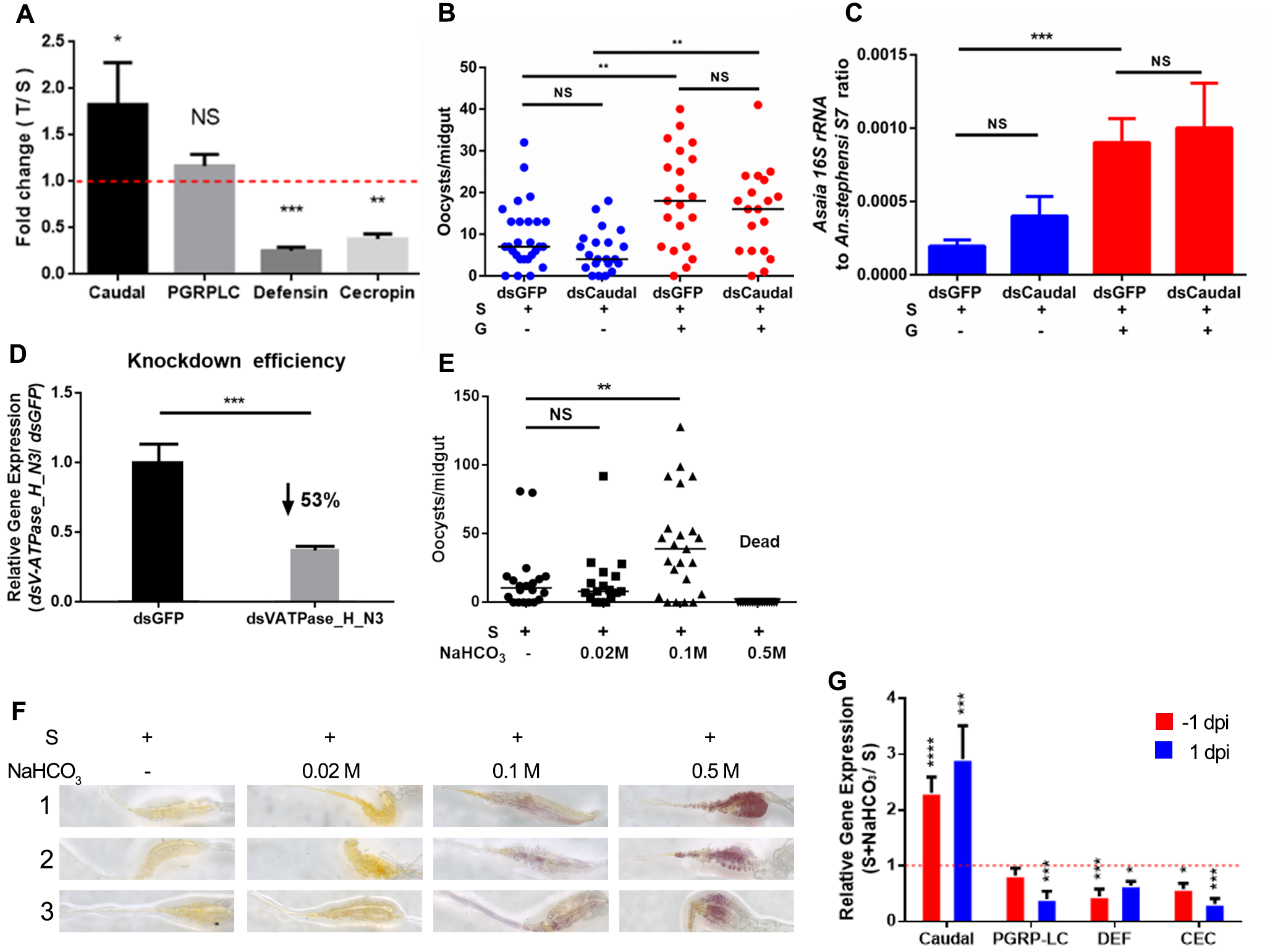

**Figure S3. Influence of mosquito midgut pH on *P. berghei* infection. Related to figure 4.**

(A) The relative expression level of *caudal, pgrp-lc, defensin,* and *cecropin* in mosquitoes fed with T -1 dpi. The expression level of genes in T feeding mosquitoes were normalized to the gene’s expression in S feeding ones.

(B) *P. berghei* oocyst intensity in dsRNA treated mosquito fed with S and G, respectively.

(C) The abundance of *A. bogorensis* in the midgut of S and G feeding *An. stephensi* treated with dsRNA.

(D) *V-ATPase_H_N3* knocking down efficiency. Relative expression level of *V-ATPase_H_N3* was normalized to that in dsGFP control. Ribosomal gene *s7* used as an internal control. Error bars indicate standard error (*n* =10). Results from one of three independent experiments are shown.

(E) *P. berghei* oocyst intensity in mosquito fed with S and S+0.02 M, 0.1 M and 0.5 M NaHCO_3_, respectively.

(F) The pH staining of mosquito midguts fed with increasing concentration of NaHCO_3_ by m-cresol purple. Images are the three representatives of at least 5 individual mosquito midguts.

(G) The expression levels of *caudal, pgrp-lc, defensin,* and *cecropin* in mosquito fed with S + 0.1 M NaHCO_3_ diet 1 day prior to (-1 dpi) and 1 day (1 dpi) post infection, were normalized to the gene’s expression of the ones fed with S diet.

Each dot in B and E represents an individual mosquito and horizontal lines represent the medians. Significance in A, C, D and G was determined by Student’s *t* test. Error bars indicate standard error of the mean (A, D and G, *n*=8, C, *n*=10). Results shown in B and E were pooled from at least 2 independent experiments. Significance was determined by Mann-Whitney test. NS, no significance, *, p <0.05, **, p<0.01, ***, p<0.001and ****, p<0.0001. T, 2% sucrose+0.1 M trehalose; S, 2% sucrose; G, 2% sucrose+0.1 M glucose.

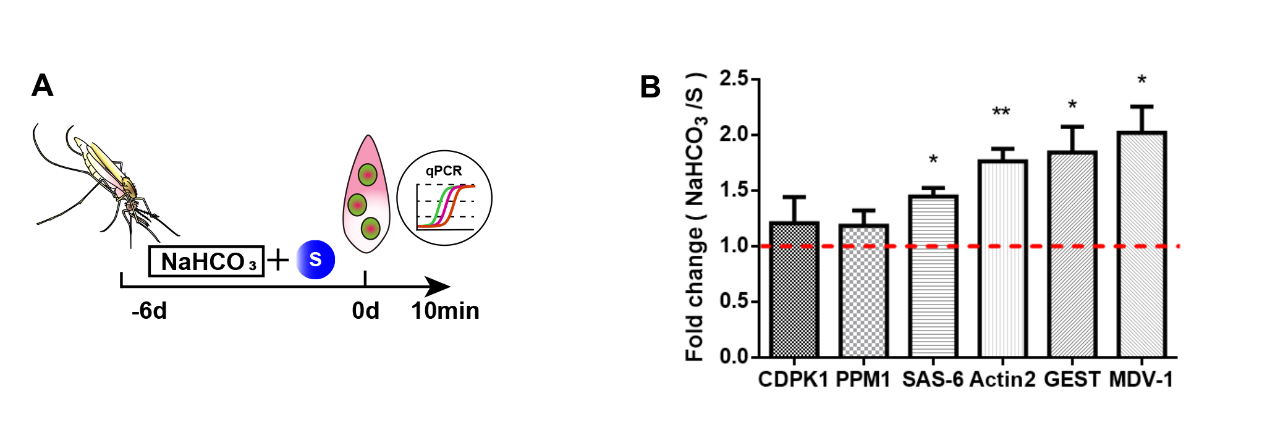

**Figure S4. Addition of NaCHO_3_ induced the expression of genes associated with microgamete development. Related to figure 6.**

(A) Work flow of NaCHO_3_ treatment and qPCR assay.

(B) Quantification of expression level of genes associated with male gametogenesis. The expression level of male gametogenesis related genes in NaHCO_3_ (0.1 M) supplemented mosquitoes was normalized to that in controls.

Significance in B was determined by Student’s *t* test. Error bars indicate standard error of the mean (*n* = 8). *, p <0.05, and **, p<0.01. S, 2% sucrose.
