## Supplemental Table 1,3,4 for "Glucose-mediated expansion of a gut commensal bacterium promotes *Plasmodium* infection through alkalizing mosquito midgut"

**Supplemental Tables:**

**Supplementary Table 1: Significantly changed metabolites with corresponding correlation coefficients in the mosquito extract between NB and IB Group.** **Related to Figure 1.** Lipids: R-C*H*_3_, R-C*H*_2_, C*H*_2_CH_2_CO, C*H*_2_C=C, UFA (unsaturated fatty acids); dimethylamine: DMA; phosphorylcholine: PC; glycerophosphocholine: GPC; Inosine 5'-phosphate: IMP; adenosine diphosphate: ADP.

|  | Keys | Metabolite | NB1D vs. IB1D |
| --- | --- | --- | --- |
| Lipids | 1 | R-C*H*_3_^a^ | -0.98 |
|  | 2 | R-C*H*_2_ | -0.99 |
|  | 3 | C*H*_2_CH_2_CO | -0.97 |
|  | 4 | C*H*_2_C=C | -0.92 |
|  | 5 | UFA | -0.97 |
| Glucose & TCA Cycles | 6 | trehalose | -0.85 |
|  | 7 | glucose | -0.88 |
|  | 8 | pyruvate | 0.60 |
|  | 9 | succinate | -0.87 |
|  | 10 | citrate | -0.77 |
|  | 11 | acetate | 0.85 |
|  | 12 | acetoacetate | 0.79 |
| Amino acids | 13 | tyrosine | 0.69 |
| Cholines | 14 | choline | 0.74 |
|  | 15 | PC | 0.79 |
|  | 16 | GPC | -0.96 |
|  | 17 | DMA | 0.60 |
| Nucleotides & Nucleosides | 18 | xanthurate | 0.74 |
|  | 19 | inosine | -0.66 |
|  | 20 | ADP | -0.88 |
|  | 21 | IMP | 0.61 |

**Supplementary Table 3: Significantly changed metabolites in the conditioned medium extracts of *An. stephensi* fed on S, G and T diets.** **Related to Figure 5.** Adenosine diphosphate, ADP, *, P<0.05, **, P<0.01.

| Keys | Metabolites | Fold change (G/S) | Fold change (T/S) |
| --- | --- | --- | --- |
| 1 | glucose | 1.24 | 0.41 ^*^ |
| 2 | citrate | 0.52 ^**^ | 0.31 ^*^ |
| 3 | succinate | 0.53 ^**^ | 0.33 ^*^ |
| 4 | fumarate | 0.70 | 0.22 ^*^ |
| 5 | lactate | 0.47 ^**^ | 0.82 |
| 6 | valine | 0.45 ^**^ | 0.74 |
| 7 | isoleucine | 0.45 ^**^ | 0.82 |
| 8 | leucine | 0.46 ^**^ | 0.82 |
| 9 | alanine | 0.39 ^**^ | 0.50 ^*^ |
| 10 | tyrosine | 0.47 ^**^ | 0.75 |
| 11 | phenylalanine | 0.45 ^**^ | 0.81 |
| 12 | tryptophan | 0.46 ^**^ | 0.76 |
| 13 | hypoxanthine | 0.33 ^**^ | 0.76 |
| 14 | uracil | 0.49 ^**^ | 0.90 |
| 15 | ADP | 0.12 ^**^ | 0.06 ^**^ |

**Supplementary Table 4: qRT-PCR primers used in this study. Related to Figure 3, Figure 6, Figure 7, Figure S4, Figure S5, Figure S6, Figure S7 and Figure S9.**

| Gene | Forward primer | Reverse primer | Reference |
| --- | --- | --- | --- |
| 16S-EUB | AGAGTTTGATCCTGGCTCAG | GGTTACCTTGTTACGACTT | (Marchesi et al., 1998) |
| VATPase_H_N3 | TGGCATCAGCACGCTCATCAG | TCCTTCGCACAGTCGCTCAGA | This paper |
| Caudal | CAAGGACCGCAAGCAGAAGAA | CGATTGACCGCCGAGACC | This paper |
| *A. bogorensis* 16S rRNA | GATGACATGAACCGTGCCCTGG | ACCTCCGTCTTGATGGCGTACA | (Favia et al., 2007) |
| *An. stephensi* S7 | TGCGGAGCGTCGTATTCTGC | ACACAGCGGTGAGCGTTCG | (Salazar et al., 1993) |
| *P.berghei* CDPK1 | TGGTGGGCAAAACGATCAAGA | GCTTCCTCAGCCGTACATCT | This paper |
| *P.berghei* PPM1 | AGGGGATAGTCGCTGTGTCTT | ACCTCGGCATACTCCTAAGCAT | This paper |
| *P.berghei* Acint2 | TTCTTGATAGTGGTGATGGCGTAA | CGATTTCTCTTTCAGCGGTTGTAG | This paper |
| *P.berghei* GEST | CTACATTCGAGAGCCATGATACTT | AAACTGTGTCACCAATTTCAAGAT | This paper |
| *P.berghei* MDV-1 | TCCAACATCAACCATAGGGTGTCT | TGCCTTGCCTCCACTTCCA | This paper |
| *P.berghei* SAS-6 | AAAGGATATGGGCGTGATGGAT | ACACTCATTAGATGTACCACCACT | This paper |
| *P.berghei* 18S | AAGCATTAAATAAAGCGAATACATCCTTAC | GGAGATTGGTTTTGACGTTTATGTG | (Baptista et al., 2010) |
| *An. stephensi* Cecropin | GCTGCTCTTTCTCGTTGCG | CGGCACCTTCCACCTTCT | This paper |
| *An. stephensi* Defensin | CCGCCTTGAACACGCTCCT | GCTGCCGACACCGAATCCA | This paper |
| *An. stephensi* PGRP-LC | TGTGCCATCGTAGCGGTCAT | AGCCACTCGGTTCTCGTCAC | This paper |
